# AAV-miR-124 enhances endogenous alveolar epithelial regenerative plasticity and reverses bleomycin-induced pulmonary fibrosis

**DOI:** 10.64898/2026.08.03.741463

**Authors:** Maria Concetta Volpe, Giulia Zandomenego, Alberto Maria Davide Ingo, Raffaella Klima, Martina Torresi, Lorena Zentilin, Paola Confalonieri, Francesco Salton, Danilo Licastro, Marco Confalonieri, Luca Braga

## Abstract

Idiopathic pulmonary fibrosis (IPF) is a progressive interstitial lung disease characterized by irreversible destruction of the alveolar epithelium and impaired regeneration. Although current therapies slow disease progression, they do not restore functional alveoli, highlighting the need for regenerative approaches that promote endogenous lung repair. Here, we performed the first unbiased functional screen of 2,042 human microRNA mimics in primary mouse alveolar type II (ATII) cells to identify regulators of ATII-to-alveolar type I (ATI) cell transdifferentiation. The screen identified miR-124-3p as the most effective promoter of ATI differentiation. In vitro, miR-124-3p promoted ATII-to-ATI transdifferentiation in healthy and bleomycin-injured ATII cells while also increasing the ATII cell pool, consistent with activity on epithelial progenitors. Using the engineered AAV6.2FF capsid, we generated a vector encoding miR-124-3p, which efficiently transduced ATII cells, MHC-II⁺ club distal progenitor cells, and injury-induced KRT8⁺ epithelial intermediates. Therapeutic administration after fibrosis establishment reduced lung fibrosis, restored alveolar architecture, and showed greater efficacy than nintedanib in the bleomycin mouse model. Mechanistically, we propose a context-dependent model whereby miR-124-3p regulates epithelial cell states through the EZH2-C/EBPα axis while attenuating epithelial transcriptional programs associated with IPF. Together, these findings support AAV-mediated delivery of miR-124 to promote alveolar repair in pulmonary fibrosis.

## Introduction

In the adult lung, alveolar type II (ATII) cells function as facultative stem cells capable of self-renewal and transdifferentiation into alveolar type I (ATI) cells, which constitute most of the gas-exchange surface ^1,2^. Failure of the ATII-to-ATI transdifferentiation program is now considered a hallmark of idiopathic pulmonary fibrosis (IPF) ^3^. IPF is a chronic, progressive interstitial lung disease of unknown aetiology, characterized by excessive fibrotic scarring of the lung parenchyma, leading to destruction of normal alveolar architecture, loss of functional alveoli, including ATII and ATI cells, and irreversible impairment of gas exchange, resulting in reduced blood oxygenation ^4,5^.

Although fibroblast activation and extracellular matrix deposition are the final pathological effectors of fibrosis, IPF is increasingly understood as a disease in which repeated epithelial injury, ageing-associated loss of regenerative competence, and senescence-related epithelial dysfunction converge to impair effective alveolar repair and regeneration by disrupting ATII-to-ATI transdifferentiation. This failure of epithelial regeneration sustains a pro-fibrotic microenvironment characterized by fibroblast activation, inflammatory crosstalk, and excessive extracellular matrix deposition.^6–8^

Recent lineage-resolved studies have refined the cellular hierarchy governing alveolar regeneration. Employing genetically engineered mouse models enabling parallel lineage tracing of distinct epithelial populations, Liu et al. demonstrated that alveolar repair is supported by mature surfactant protein C–expressing (SPC⁺) ATII cells, with additional contributions from bronchioalveolar stem cells (BASCs) ^9–11^ and airway-derived club cells, including H2 K1⁺ and MHC-II⁺ Club distal progenitor cells ^8,11,12^, which participate in distal alveolar regeneration by adopting an ATII cell fate and subsequently differentiating into ATI cells. These progenitors converge on a KRT8⁺ transitional epithelial state that is required for effective regeneration but becomes pathogenic when persistently maintained by fibroblast- and macrophage-driven signalling ^8,13^.

Together, these findings have shifted the conceptual framework of IPF from a disorder primarily driven by fibroblast dysfunction to one initiated and sustained by epithelial failure ^14,15^. Consequently, therapeutic strategies aimed at restoring physiological alveolar epithelial repair mechanisms represent a major unmet need. However, the rational design of such regenerative approaches remains challenging due to the lack of discrete, targetable regulators of epithelial plasticity and the difficulty of reducing complex pathological states to isolated molecular functions. For this reason, therapeutic modulation of epithelial plasticity may therefore require molecules with inherently pleiotropic activity, capable of acting through coordinated modulation of multiple targets rather than single, fully defined interactions.

MicroRNAs (miRNAs) are small non-coding RNAs that regulate gene expression post-transcriptionally and, by simultaneously modulating multiple targets, exert intrinsically pleiotropic effects that are central to epithelial differentiation, cell fate decisions, and tissue regeneration ^16^. In the lung, numerous microRNAs have been implicated in IPF and fibrotic remodelling, primarily through regulation of fibroblast activation or epithelial-to-mesenchymal transition, including miR-let7d ^17^, miR-29 ^18,19^, miR-21 ^20,21^, miR-200 ^22^, miR-26a ^23^, miR-503 ^24^, and miR-155 ^25^. To date, only miR-375 ^26^ and miR-200 ^27,28^ have been associated with ATII-to-ATI transdifferentiation and alveolar repair.

More recently, Chioccioli et al. demonstrated systemic delivery of a peptide-conjugated miR-29 mimic to lung fibroblasts, achieving antifibrotic efficacy and a favourable safety profile in rodent and non-human primate models ^29^. Despite this progress, systemic microRNA mimic delivery remains limited by poor stability, rapid degradation, restricted tissue penetration, and limited cell-type specificity. Viral vectors, by contrast, offer superior tissue and cellular tropism and enable efficient gene transfer in vivo, although their clinical application requires careful consideration of prolonged transgene expression and manufacturing complexity.

Among available platforms, adeno-associated virus (AAV) vectors are particularly well suited for epithelial-directed delivery. AAVs represent a clinically validated gene-delivery system, with over 238 clinical trials employing recombinant AAV-based therapies and seven approved products ^30^. The AAV6.2FF variant, derived from naturally occurring AAV6 ^31,32^, exhibits enhanced transgene expression in the distal lung epithelium with comparatively minimal fibroblast transduction and has demonstrated therapeutic efficacy in preclinical lung disease models ^33,34^. Nevertheless, despite the availability of efficient lung epithelial gene-delivery platforms, no systematic effort has yet been undertaken to identify microRNAs that directly regulate ATII-to-ATI trans-differentiation.

In this study, we performed high-throughput screening (2042 human microRNA mimics) in primary mouse ATII cells to identify microRNAs that promote this process, leading to the identification of miR-124-3p and miR-5008-5p as candidate regulator. We established an AAV6.2FF-microRNA expression system to achieve efficient overexpression of individual microRNA precursors in alveolar epithelial cells in vivo and evaluated their therapeutic potential in the bleomycin-induced mouse model of lung fibrosis ^35^. To directly assess epithelial regeneration in vivo, we employed SFTPC-Cre–based lineage tracing ^2^, we demonstrate miR-124-3p-induced ATII-to-ATI transdifferentiation and we showed that AAV6.2FF transduces mature ATII cells, MHC-II⁺ Club distal progenitor cells, and injury-induced KRT8⁺ epithelial intermediates, enabling modulation of epithelial programs that allow fibrotic tissue remodelling by leveraging effective alveolar repair.

Consistent with this regenerative interpretation, we found that endogenous miR-124-3p levels increased following acute bleomycin-induced lung injury in young mice, peaking at day 21 before declining during spontaneous resolution, whereas this induction was largely absent in aged mice ^36^, which exhibit impaired spontaneous injury resolution, suggesting that miR-124-3p may participate in physiological alveolar repair programmes that are attenuated with ageing.

Building on these findings, together with the previously described role of miR-124-3p in embryonic alveologenesis ^37^ and the concept that alveolar repair after injury and developmental alveologenesis rely on partially shared molecular programs ^38^, we focused on two established miR-124-3p targets, EZH2 and C/EBPα, recently implicated in regulating ATII cell plasticity during lung development, repair after injury^39^, and fibrosis-associated epithelial remodelling ^40^.

We further demonstrate that miR-124-3p operates in a context-dependent manner by engaging distinct targets across different epithelial cell states. In MHC-II⁺ Club distal progenitor cells, miR-124-3p targets EZH2, thereby facilitating epithelial maturation programs and stabilization of ATII cell identity, whereas in mature ATII cells miR-124-3p promotes reparative plasticity and ATII-to-ATI transdifferentiation through modulation of C/EBPα-dependent transcriptional programs. These findings suggest that miR-124-3p does not enforce a single epithelial fate but rather coordinates distinct regenerative programs across multiple stages of the alveolar epithelial hierarchy.

## Results

### Functional Screening of 2,042 Human microRNA Mimics in Primary Mouse ATII Cells

First, we performed a cell-based phenotypic high-throughput screening (HTS) assay on 2,042 human microRNA mimics to evaluate their efficacy in promoting ATII-to-ATI transdifferentiation in primary alveolar mouse epithelial cells. To our knowledge, this represents the first arrayed functional screening of mature microRNA mimics in primary mouse ATII cells. To achieve this, we developed an *in vitro* assay leveraging the natural tendency of ATII cells to transdifferentiate into ATI cells when cultured in 2D ^1^. This approach builds on similar assays used in our previous studies ^27,28^ to investigate the mechanisms regulating ATII-to-ATI transdifferentiation. For this study, we adapted the assay to be compatible with an automated pipeline for image-based functional screenings. We isolated primary ATII cells from 8-week-old C57BL/6 mice as described in ^28^ and reverse-transfected them with an arrayed library of 2,042 human microRNA mimics at a final concentration of 25nM (Horizon Discovery, miRBase V.19). As shown in Supplementary Fig. 1A-G, four days after seeding, an average of 40% of primary mouse ATII cells spontaneously transdifferentiated into ATI cells, stained for RAGE. Based on this observation, we selected this time point to assess microRNA effects, enabling the identification of both inhibitors and promoters of transdifferentiation.

At four days post-transfection, cells were fixed and immunostained for RAGE, all wells were imaged, and the images were processed using automated image analysis to quantify the total number of cells and RAGE+ (ATI) cells. Plates were next immunostained for p21, a marker of cellular senescence, in order to further characterize microRNAs that were blocking ATII-to-ATI transdifferentiation by inducing cellular senescence (data not shown).

First, we excluded all toxic microRNAs, defined as those with a Z-score for the total cell count ≤ -1.96 (p ≤ 0.05) (Fig. 1B). Next, we ranked the remaining microRNAs based on their ability to either increase (Z-score for ATI cells per well ≥ 1.96, p ≤ 0.05) or decrease (Z-score for ATI cells per well ≤ -1.96, p ≤ 0.05) the number of ATI cells per well (Fig. 1C).

**Figure 1.**
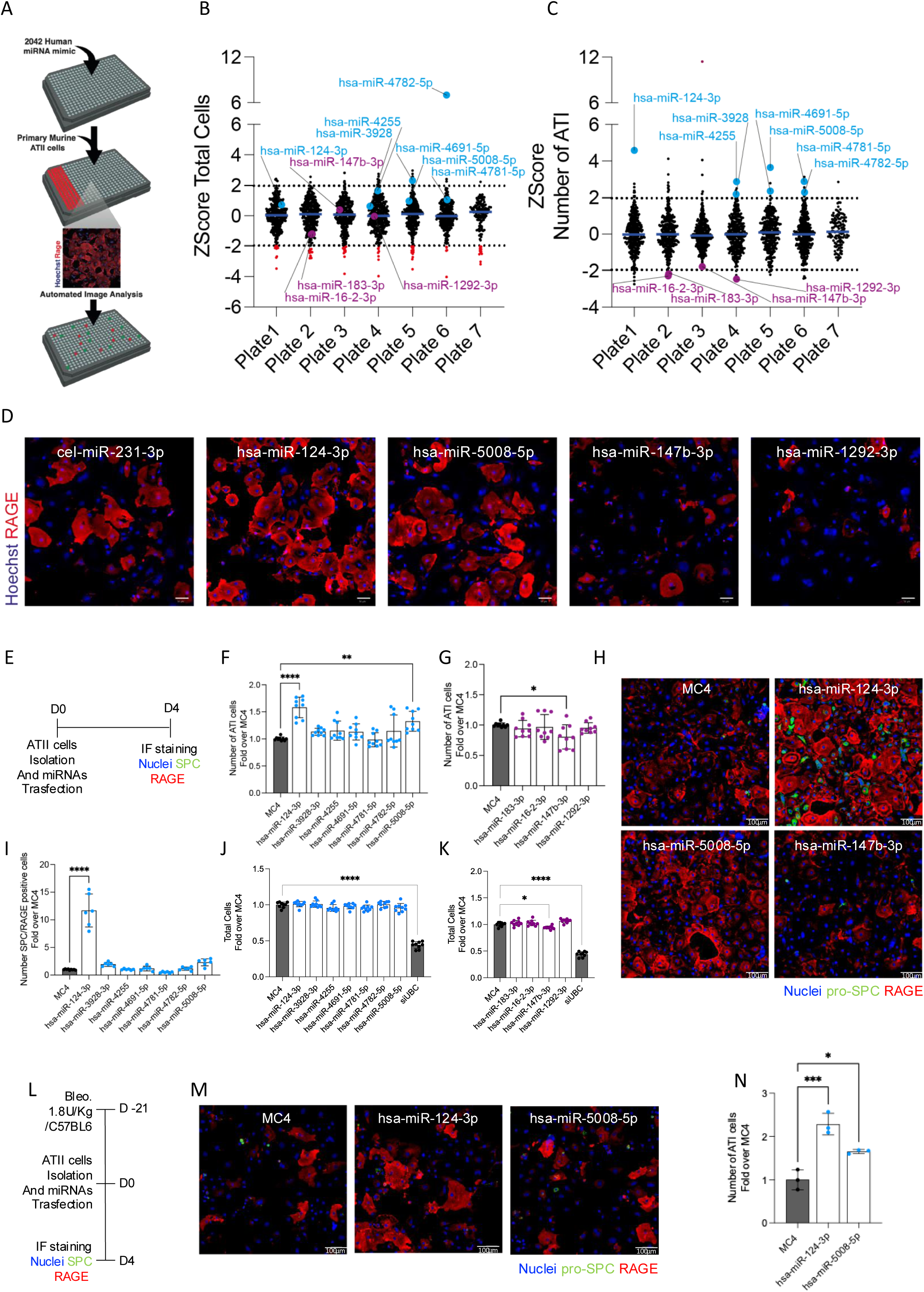
Cell-based phenotypic high-throughput screening (HTS) of human miRNA mimics in primary murine ATII cells. (A) Schematic overview of the HTS workflow. A library of 2,042 human miRNA mimics was reverse-transfected (final concentration 25 nM) into primary murine alveolar type II (ATII) cells. (B) Z-score analysis of total cell number, used to discriminate toxic (red) from non-toxic miRNAs. (C) Z-score of ATII cell number normalized to the negative control (cel-miR-231-3p), identifying miRNAs that promote ATII-to-ATI transdifferentiation (upper panel, light blue) and miRNAs that inhibit ATII-to-ATI transdifferentiation (lower panel, violet). (D) Representative immunofluorescence images of miRNAs promoting ATII-to-ATI transdifferentiation (hsa-miR-124-3p and hsa-miR-5008-5p) and miRNAs inhibiting transdifferentiation (hsa-miR-147b-3p and hsa-miR-1292-3p); ATI cells are stained for RAGE (red), and nuclei are counterstained with Hoechst (blue). (E) Schematic representation of the validation strategy. Primary murine ATII cells were isolated and transfected with selected miRNAs (25 nM final concentration). (F–G) ATII cell number, expressed as fold change over the negative transfection control (MC4), for miRNAs promoting (F) or inhibiting (G) ATII-to-ATI transdifferentiation (three independent experiments). (H) Representative immunofluorescence images of primary murine ATII cells transfected with MC4 (control), miRNAs that significantly increased ATI cell number (hsa-miR-124-3p and hsa-miR-5008-5p), or decreased ATI cell number (hsa-miR-147b-3p); ATI cells are stained as in (D), with additional staining for pro-SPC (green). (I) Quantification of transitional ATII cells (pro-SPC⁺/RAGE⁺), expressed as fold change over MC4. (J–K) Evaluation of miRNA toxicity, expressed as total cell number relative to MC4. (L) Schematic overview of miRNA validation in bleomycin-treated ATII cells: mice were intratracheally injected with bleomycin (1.8 U/kg), and after 21 days ATII cells were isolated and transfected with hsa-miR-124-3p and hsa-miR-5008-5p; cells were fixed after 4 days and stained as in (H). (M) Representative immunofluorescence images of primary murine bleomycin-treated ATII cells transfected with MC4 (control), hsa-miR-124-3p, hsa-miR-5008-5p, or their combination; ATI cells are stained as in (H). (N) Quantification of total ATI cell number, expressed as fold change over MC4 across the experimental conditions shown in (M). Statistical significance was determined by one-way ANOVA followed by Dunnett’s multiple comparisons test. *P < 0.05; **P < 0.01; ***P < 0.0001.

Using this approach, we identified 47 microRNA mimics that significantly promoted ATII-to-ATI transdifferentiation and 20 microRNA mimics that significantly inhibited the process. We then proceeded with image inspection of all selected microRNAs thus selecting 7 human microRNAs as the most effective in promoting ATII-to-ATI transdifferentiation (light blue dots in Fig. 1B and Fig. 1C) and 4 human microRNAs as the most effective in blocking ATII-to-ATI transdifferentiation (Violet dots in Fig. 1B and Fig. 1C). Representative IF images for selected microRNAs are reported in Fig. 1D.

### Secondary screening and validation of pro-differentiating microRNA in diseased ATII cells

Given the naturally limited yield of ATII cells from adult mouse lungs, the primary screening was not conducted in duplicate. Therefore, a secondary screening aimed at validating the selected microRNA mimics represents an essential step. In this phase of the study, we focused on addressing three major questions: (1) confirming efficacy on primary ATII cells; (2) excluding unwanted pro-fibrotic effects on lung fibroblasts; and (3) excluding potential pro-tumorigenic effects, considering the relevance of ATII cells in lung adenocarcinoma development ^41^.

Validation was performed for all 11 selected microRNAs and on three independent isolations of ATII cells by using the same transdifferentiation assay used for the primary screening but in 96 well plates and with all microRNA mimics newly procured from Horizon Discovery.

Following isolation, cells were reverse transfected with selected microRNA mimics at the final concentration of 25 nM. As a negative control, cel-miR-231-3p (non-targeting MC4) was used, while siUBC, known to induce reduced viability upon effective transfection, served as the positive control. The assay was terminated 4 days post-transfection, after which cells were fixed and immunostained for RAGE (ATI cells) and SPC (ATII cells), with nuclei counterstained using Hoechst 33342 (experimental scheme shown in Fig. 1E).

The positive control siUBC effectively reduced cell viability upon transfection (Fig. 1J,K), thereby confirming transfection efficiency. Among the seven selected pro-differentiation microRNAs, hsa-miR-124-3p and hsa-miR-5008-5p proved to be the most effective in increasing the number of ATI cells per well, suggesting enhanced ATII-to-ATI transdifferentiation (Fig. 1F). In contrast, as shown in Fig. 1G, hsa-miR-147b-3p resulted in the most substantial reduction in ATI cell number, accompanied by a significant loss in viability, thus leaving its actual effect on negatively regulating ATII-to-ATI transdifferentiation debatable.

Representative images of all treatments (Fig. 1H) clearly show that treatment with hsa-miR-124-3p not only led to an increased number of ATI cells three days post-treatment, but also to a marked rise in both SPC-positive and SPC and RAGE double-positive cells (Fig. 1I).

Subsequently, we evaluated all 11 microRNAs in mouse fibroblasts stimulated with TGF-β. None of the tested microRNAs enhanced TGF-β-induced myofibroblast transition or promoted myofibroblast proliferation (Supplementary Fig. 2A-C). Notably, hsa-miR-124-3p did not counteract the effects of TGF-β, but significantly reduced the proliferation of stimulated myofibroblasts.

Regarding safety, none of the 11 microRNAs induced proliferation of the A549 adenocarcinoma cell line, with hsa-miR-124-3p once again emerging as the most potent anti-proliferative candidate (Supplementary Fig. 2D-F). These results are consistent with current literature on miR-124-3p in cancer, where it has been described as a tumour suppressor across multiple cancer types, including Non-Small Cell Lung Cancer (NSCLC), by inhibiting proliferation, migration, and metastasis ^42–44^. In NSCLC, it has been shown to negatively regulate EMT by targeting CDH2 ^44^, and similarly in triple-negative breast cancer, by targeting ZEB2 ^45^. Of note, both ZEB1 and ZEB2 are well-established regulators of EMT also in idiopathic pulmonary fibrosis ^46–48^.

Lastly, we tested the efficacy of hsa-miR-124-3p, hsa-miR-5008-5p on ATII cells isolated from bleomycin-treated mice at 21 days post-administration. For this purpose, we applied the same differentiation assay used in the primary and secondary screenings (Fig. 1L). As shown by the representative images in Fig. 1M and the quantifications in Fig. 1N, both microRNAs confirmed their ability to promote ATII-to-ATI transdifferentiation in diseased ATII cells *in vitro,* with hsa-miR-124-3p emerging as the most effective.

These results highlight hsa-miR-124-3p and hsa-miR-5008-5p as the two most effective microRNAs in promoting ATII-to-ATI transdifferentiation in both primary healthy and diseased mouse ATII cells, with miR-124-3p being the most effective of the two. In addition, miR-124-3p was the only one to also increase the number of ATII cells and double positive SPC/RAGE intermediate cells, suggesting its potential efficacy not only on mature ATII cells but also on ATII cell progenitors. Neither of the two selected microRNAs showed any unwanted secondary effects on primary lung fibroblasts or A549 cells, with miR-124-3p displaying a strong anti-proliferative effect, which supports its safety and which is consistent with its observed pro-differentiation activity on ATII cells.

### *In vivo* microRNA delivery to alveolar epithelial cells using engineered pseudo-typed AAV vector

Given the original hypothesis that the selected microRNAs could enhance endogenous alveolar repair mechanisms by promoting ATII-to-ATI transdifferentiation, we opted for an *in vivo* delivery strategy designed to preferentially express microRNA into alveolar epithelial cells. To this end, we employed adeno-associated viral (AAV) vectors, specifically using an artificially engineered capsid variant, AAV6.2FF ^31,32^.

As an initial step, we validated the delivery strategy by administering 3 × 10¹¹ vg/mouse of an AAV6.2FF-GFP vector to healthy 8-week-old C57BL/6 mice via intratracheal instillation. Lung transduction was assessed at days 5, 10, 21, 30, and 60 post-injection (experimental scheme in Fig. 2A). GFP expression, measured by qRT-PCR on total lung lysates, was robust at all time points and peaked at day 10 (Fig. 2B). This kinetic profile was mirrored at the cellular level, where analysis of GFP⁺/SPC⁺ alveolar type II (ATII) cells revealed a peak of transduction at day 5 (35.89% ± 8.8%), gradually declining to 15.88% ± 2.3% by day 60 (Fig.s 2C and 2D). As expected, no GFP-positive fibroblasts were detected by immunofluorescence staining (data not shown), consistent with the preferential specificity AAV6.2FF serotype for lung epithelial cells ^32,33^. These results were further supported by flow cytometry, confirming an average of 23.83% ± 6.3% of GFP⁺ SPC+ cells (Fig.s 2E and 2F).

**Figure 2.**
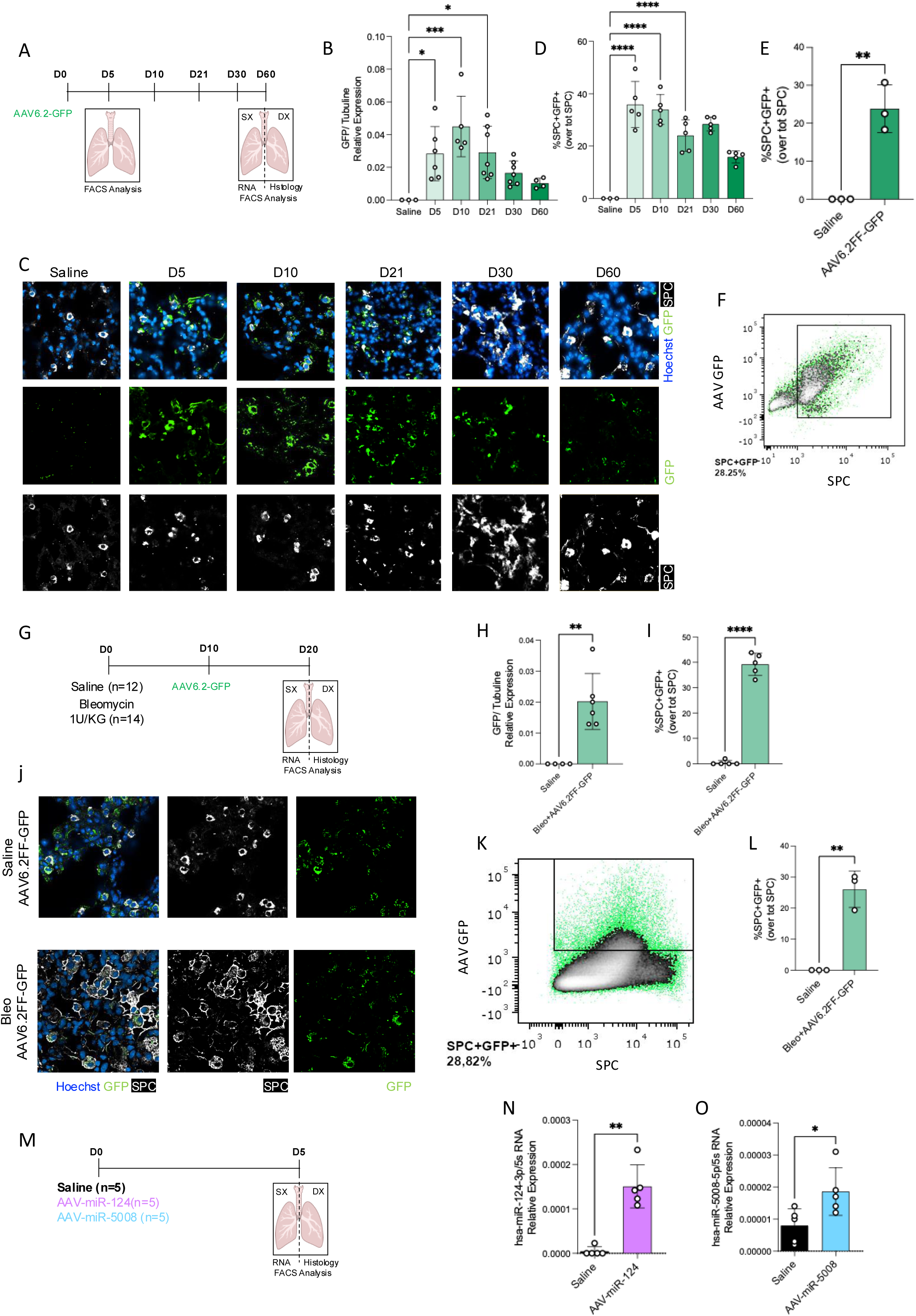
Kinetics and transduction efficiency of AAV6.2FF in healthy and bleomycin-treated mice. (A) Schematic overview of the experimental workflow. AAV6.2FF-GFP (30μL, 1 × 10¹³ vg/mL) was administered by intratracheal injection to 34 mice. Animals were sacrificed at 5, 10, 21, 30, and 60 days post-injection. Lungs were harvested, with the right lobe processed for RNA extraction and flow cytometry analysis and the left lobe used for histological analysis (immunofluorescence staining). (B) Quantification of GFP mRNA expression in the right lung lobe by real-time PCR, normalized to tubulin. (C) Representative immunofluorescence images of lung sections collected at 5, 10, 21, 30, and 60 days after intratracheal administration of AAV6.2FF-GFP or saline (control), with ATII cells stained for pro-SPC (white), GFP in green, and nuclei counterstained with Hoechst (blue). (D) Quantitative image analysis of AAV6.2FF-GFP–transduced ATII cells by immunofluorescence. (E) Flow cytometry quantification of transduced ATII cells at 5 days post-injection, showing that approximately 24% of ATII cells were GFP-positive. (F) Representative schematic of the flow cytometry gating strategy used to identify ATII cells (pro-SPC– positive population) and GFP-positive cells. (G) Schematic overview of the experimental design for AAV6.2FF transduction in bleomycin-treated mice. Bleomycin (1.8 U/kg) was administered by intratracheal injection; ten days later, mice received intratracheal AAV6.2FF-GFP and were sacrificed 10 days after AAV administration. The right lung lobe was used for RNA extraction and flow cytometry, and the left lobe for immunofluorescence analysis. (H) Quantification of GFP mRNA expression in the right lung lobe by real-time PCR, normalized to tubulin. (I) Quantitative image analysis of AAV6.2FF-GFP–transduced ATII cells in bleomycin-treated mice, showing that approximately 39% of pro-SPC⁺ cells were GFP-positive. (J) Representative immunofluorescence images of lung sections collected 21 days after bleomycin injection (10 days after AAV6.2FF-GFP administration), with ATII cells stained for pro-SPC (white), GFP in green, and nuclei counterstained with Hoechst (blue). (K) Representative schematic of the flow cytometry gating strategy for identification of ATII (pro-SPC–positive) and GFP-positive cells. (L) Flow cytometry quantification of AAV6.2FF-GFP–transduced ATII cells in bleomycin-treated mice, showing approximately 26% GFP-positive ATII cells. (M) Schematic overview of the experimental design to evaluate the efficiency of AAV6.2FF-mediated delivery of miR-124 and miR-5008 and to assess strand specificity (3p vs 5p). Mice were intratracheally injected with 30μL of AAV6.2FF-miRNA vectors (1 × 10¹³ vg/mL), and lungs were harvested 7 days post-injection for RNA analysis. (N–O) LNA-based quantitative PCR analysis of miR-124-3p (N) and miR-5008-5p (O) expression in total lung tissue 7 days after intratracheal AAV6.2FF administration. Statistical significance for multiple comparisons was determined using one-way ANOVA followed by Dunnett’s multiple comparisons test, whereas pairwise comparisons were analyzed using an unpaired two-tailed Welch’s t-test.. *P < 0.05; **P < 0.01; ***P < 0.0001.

Given our goal of overexpressing the selected microRNAs in the bleomycin model to assess their efficacy in promoting lung repair and attenuating fibrosis, we next evaluated if bleomycin injury affects AAV transduction efficiency. Because bleomycin elicits inflammatory response in the first days after administration, that could hinder vector uptake or expression, we assessed whether AAV performance is maintained under these injury conditions.

We selected day 10 after bleomycin administration for AAV delivery, as this time point marks the resolution of the acute inflammatory phase ^49^ and represents a suitable window for therapeutic microRNA administration, given the peak of fibrotic remodelling occurring between day 14 and 21 ^49^ (experimental scheme in Fig. 2G).

As shown in Fig. 2H–J, AAV6.2FF-GFP administration 10 days after bleomycin did not reduce transduction efficiency. GFP expression levels were comparable to those of non–bleomycin mice across multiple readouts, including GFP mRNA by qRT-PCR (Fig. 2H), the proportion of GFP⁺ ATII cells (39.2% ± 4.4%) by immunofluorescence (Fig. 2I), and 26.1% ± 5.9% by flow cytometry analysis (Fig. 2K,L).

Next, we generated constructs encoding the cDNA sequences of human miR-124 and miR-5008, under the control of CMV promoter, and confirmed robust mature microRNA expression in vitro in A549 cells following plasmid transfection (Supplementary Fig. 4 A-F). We then produced AAV6.2FF viral preparations expressing the selected microRNAs and validated their *in vivo* efficacy in healthy 8-week-old C57BL/6 mice (experimental scheme in Fig. 2M). At 5 days post-intratracheal injection, we observed a clear increase in mature microRNA levels in total lung RNA, as measured by qRT-PCR using LNA-based detection (Fig. 2. N,O).

### AAV6.2FF-miR-124 confers therapeutic efficacy in bleomycin-induced lung fibrosis

Based on the strong effect of both selected microRNAs in promoting ATII-to-ATI transdifferentiation *in vitro*, in both healthy and diseased ATII cells, we next evaluated their efficacy *in vivo* using the bleomycin-induced mouse model of acute lung injury ^35^. To ensure that any observed benefit was restricted to epithelial-driven repair mechanisms, we employed AAV6.2FF viral vectors for delivery. We first aimed to assess efficacy under prophylactic conditions, and if successful, to proceed with therapeutic testing. Under prophylactic conditions, on day 0, C57BL/6 mice were anesthetized, intubated, and administered intratracheally with 1.8 U/kg of bleomycin alone (Bleomycin group), or in combination with 3×10¹¹ viral genomes of AAV6.2FF-GFP, AAV6.2FF-miR-124, or AAV-miR-5008. Mice treated with saline (0.9% NaCl) served as negative controls. To compare the efficacy of our treatment with the current standard therapy for human IPF, an additional group of bleomycin-treated animals received Nintedanib (60 mg/kg, twice daily (BID)) by oral gavage starting on day 10 (experimental scheme in Supplementary Fig. 5A). At 21 days post-bleomycin, all animals were sacrificed, lungs were harvested and weighed, with the left lobe processed for Masson’s Trichrome staining, and the remaining tissue used for hydroxyproline (HP) content measurement and bulk RNA extraction. Mice treated with AAV6.2FF-miR-124 showed improved histological outcomes on Masson’s Trichrome staining (Supplementary Fig. 5B), with higher alveolar density and reduced fibrotic remodelling. These results were further supported by a reduction in total lung weight and a marked decrease in lung HP content (Supplementary Fig. 5C,D). In contrast, AAV6.2FF-miR-5008 failed to show any beneficial effect *in vivo* and did not improve any of the measured outcomes, suggesting it is ineffective in sustaining repair mechanisms in this context and was therefore not pursued further (Supplementary Fig. 5B,C,D). The inefficacy of miR-5008-5p could be partially motivated by a lower expression of miR-5008-5p compared to miR-124-3p that on the contrary showed almost 20 fold higher expression levels at 21 days post injection, as shown in Supplementary Fig. 5 G,H.

A major challenge in the development of antifibrotic therapies is demonstrating efficacy under therapeutic conditions, as IPF patients typically present with already established and clinically evident fibrosis at the time of diagnosis. Therefore, to evaluate the translational relevance of our findings, we next assessed AAV6.2FF-miR-124 activity in a therapeutic bleomycin model, where treatment was initiated 10 days following bleomycin injury (experimental scheme in Fig. 3A). In this setting as well, miR-124-3p treatment led to improved histological appearance with increased alveolar density (Fig. 3B), and a clear reduction in fibrosis, as shown by lower lung weight and HP content (Fig. 3C, D). Notably, in both prophylactic and therapeutic protocols, miR-124-3p outperformed Nintedanib in reducing hydroxyproline content and showed a more favorable trend across all other assessed parameters, including body weight loss and overall survival (Fig. 3C,D, Supplementary Fig. 5C–F, and Supplementary Fig. 6F,G). Next, we demonstrated that, under this therapeutic condition, AAV6.2FF-miR-124 effectively induced overexpression of mature miR-124-3p. This was confirmed by miRCURY LNA microRNA-based quantitative PCR performed on total lung samples (Supplementary Fig. 6C), as well as by in situ hybridization on histological sections using a miR-124-3p–specific probe (Supplementary Fig. 6C–E). Notably, bleomycin-treated mice also displayed an increased percentage of miR-124-3p–positive cells (∼28%) compared with saline controls (∼10%), thus suggesting a physiological role of miR-124-3p in response to bleomycin induced damage.

**Figure 3.**
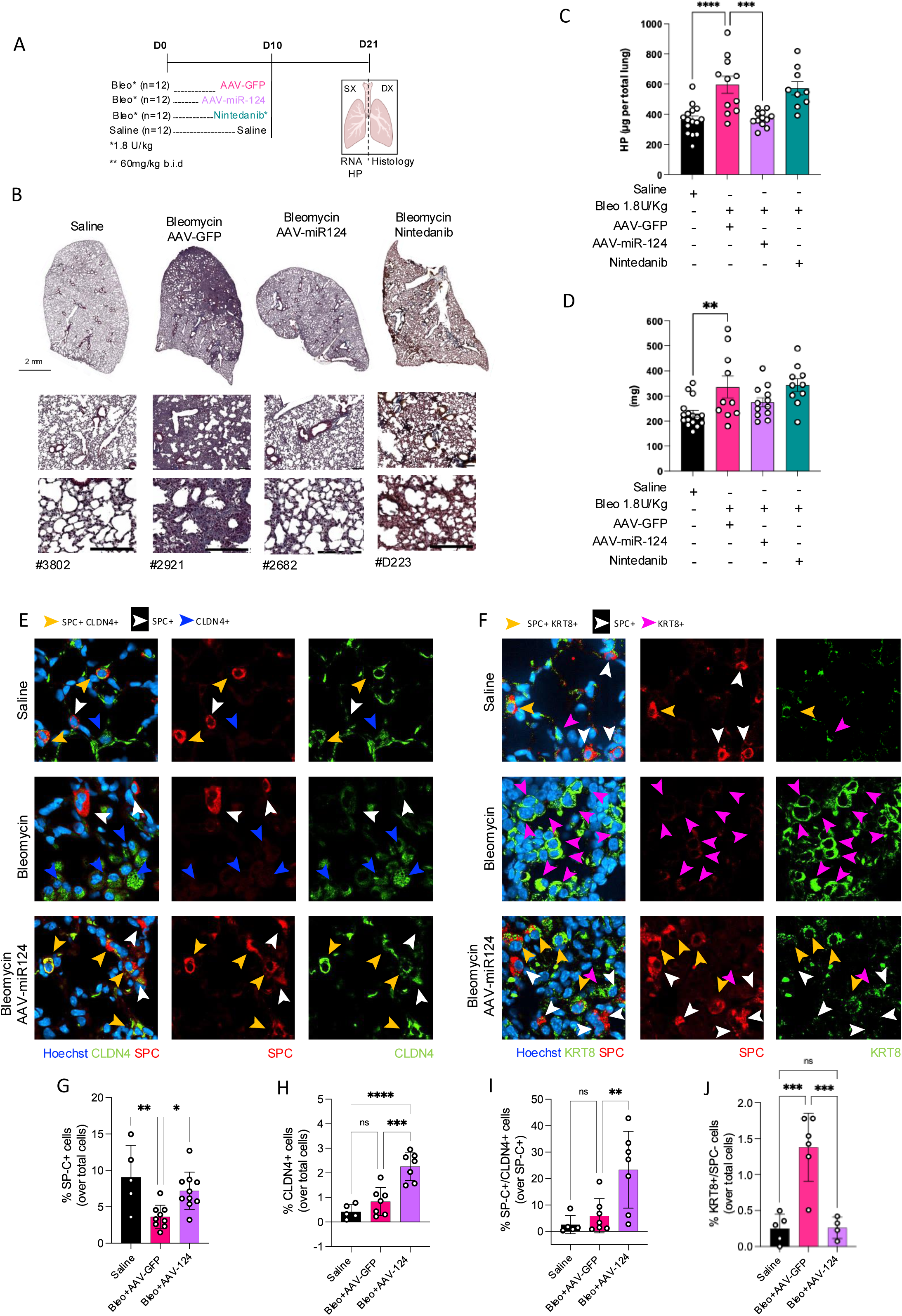
AAV6.2FF-mediated delivery of miR-124 promotes resolution of bleomycin-induced lung fibrosis. (A) Experimental scheme. Mice received intratracheal bleomycin, and 10 days later, at the onset of the fibrotic phase, were treated with AAV6.2-GFP (negative control), AAV6.2FF-miR-124, or nintedanib. Mice were sacrificed 21 days after bleomycin administration. Lungs were harvested, with the right lobe processed for RNA extraction and hydroxyproline quantification, and the left lobe used for histological analyses, including Masson’s trichrome staining and immunofluorescence. (B) Representative Masson’s trichrome staining of lung sections from mice treated as in (A). (C–D) Quantification of lung hydroxyproline content (C) and lung weight (D) across treatment groups. Representative immunofluorescence images of lung sections collected 21 days after bleomycin injection (10 days after AAV6.2FF-miR-124 administration) are shown. (E–F) ATII cells stained for pro-SPC (red) together with CLDN4 (green) (E) or KRT8 (green) (F), with nuclei counterstained with Hoechst (blue). (G–J) Quantitative image analysis showing the percentage of ATII cells (G), CLDN4-positive cells (H), CLDN4⁺/SPC⁺ double-positive transitional ATII cells (I), and KRT8⁺/SPC⁻ aberrant transitional intermediates (J) across treatment groups. Statistical significance was determined by one-way ANOVA followed by Dunnett’s multiple comparisons test. *P < 0.05; **P < 0.01; ***P < 0.0001.

Given the strong phenotypic changes sustained by miR-124-3p over-expression in therapeutic conditions, we investigated key epithelial cell states associated with alveolar repair to assess the regenerative extent of this treatment. Treatment with miR-124-3p led to a significant increase in the total number of ATII cells (Fig. 3E,F,G), indicating their preservation, restoration by acting on pre-ATII cells, or both. These findings are consistent with the results shown in Fig. 1H, where miR-124-3p was the only candidate microRNA among those tested to significantly increase the number of ATII cells in vitro. Notably, we observed an expansion of CLDN4⁺ cells (Fig. 3 E,H) and of the CLDN4⁺/SPC⁺ double-positive population (Fig. 3E,I). This aligns with findings by Zemans et al. ^13^, who identified two distinct transitional states in both human IPF and mouse models: a transitional AEC2/ABI1 (over-expressing CLDN4, KRT8, and ITGB6) state capable of progressing toward the ATI state and alveolar repair, and an aberrant basaloid/ABI2 state marked by persistent expression of basal cell genes and failure to differentiate. Accordingly, miR-124 effectively reduced the number of KRT8⁺/SPC⁻ cells at 21 days post bleomycin (Fig. 3F,J), thus further confirming its effectiveness in pushing ATII cells to assume a productive transitional state, similar to human AEC2/ABI, associated with successful alveolar regeneration while avoiding entry into aberrant, non-resolving epithelial phenotypes^8,13^.

### miR-124-3p enhances alveolar epithelial regeneration *in vitro* across distinct epithelial cell states

To further prove that miR-124-3p induces ATII-to-ATI transdifferentiation, we applied our established *in vitro* transdifferentiation protocol to ATII cells isolated from tamoxifen-treated SFTPC-CreER[T2]/mTmG mice. This genetic model enables tamoxifen-inducible lineage tracing of ATII cells by labelling all SFTPC-expressing cells with GFP. Genetically traced ATII cells were reverse-transfected with either hsa-miR-124-3p or a non-targeting MC4 control and cultured in 96-well plates for 4 days (Exp. Scheme in Fig. 4A). After fixation, cells were immunostained for RAGE: GFP+/RAGE+ cells corresponded to mature ATII cells that underwent transdifferentiation into ATI cells, whereas GFP-/RAGE+ cells represented ATI cells originating from cells that were SFTPC-negative at the time of isolation. Surprisingly, we observed minimal activity of hsa-miR-124-3p on GFP+ ATII cells *in vitro*, with no significant difference in the number of GFP+/RAGE+ cells four days after transfection (Fig. 4 B,C). Conversely, we observed a pronounced increase in GFP-/RAGE+ (Fig. 4B,D) cells upon treatment with hsa-miR-124-3p. These findings well align with previous evidence from our secondary validation screening, which indicated that hsa-miR-124-3p uniquely promotes also an increase in ATII cells and SPC/RAGE double-positive intermediate cells (Fig. 1H). This further suggests a potential hsa-miR-124-3p mechanism of action mediated through effects on ATII progenitor cells.

**Figure 4.**
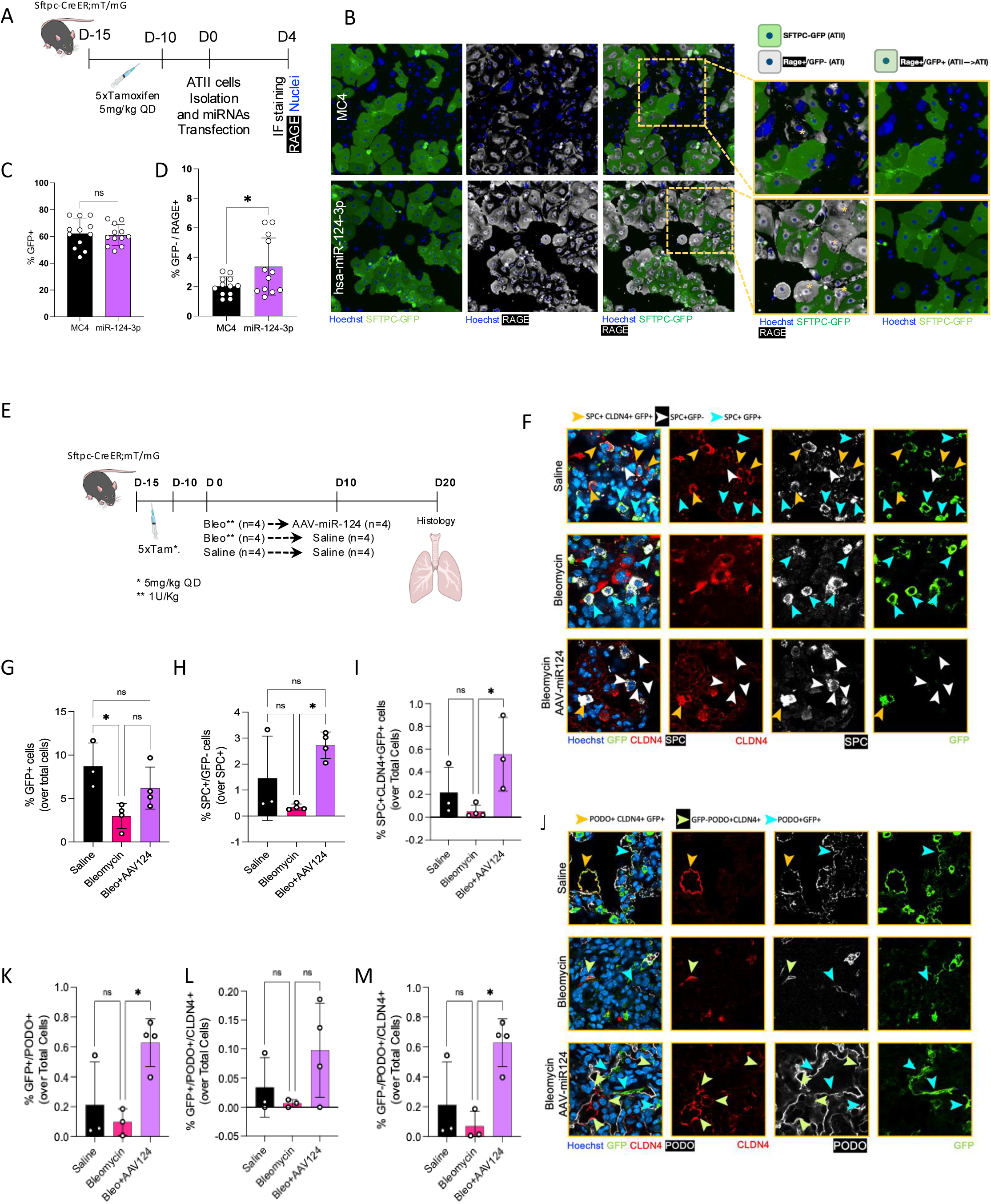
Lineage tracing analysis of miR-124-3p–mediated ATII-to-ATI transdifferentiation in vitro and in vivo. (A) Schematic overview of the in vitro lineage tracing experimental design used to assess the ability of miR-124-3p to induce ATII-to-ATI transdifferentiation. SFTPC-CreER[T2]/mTmG mice received intraperitoneal tamoxifen injections for 5 consecutive days, followed by a 10-day washout period. ATII cells were then isolated, transfected with hsa-miR-124-3p, and fixed 4 days post-transfection for analysis. (B) Representative immunofluorescence images of primary ATII cells isolated from SFTPC-CreER[T2]/mTmG mice and transfected with hsa-miR-124-3p, with ATI cells stained for RAGE (white) and nuclei counterstained with Hoechst (blue). (C–D) Quantification of GFP-positive cells and GFP⁻/RAGE⁺ cells 4 days after transfection. (E) Schematic overview of the in vivo lineage tracing experimental design in bleomycin-induced lung fibrosis. SFTPC-CreER[T2]/mTmG mice received intraperitoneal tamoxifen injections for 5 consecutive days, followed by a 10-day washout period, after which bleomycin was administered intratracheally. Ten days later, animals received intratracheal AAV6.2FF-miR-124. Mice were sacrificed 21 days after bleomycin administration, and lungs were collected for histological analysis. Representative immunofluorescence images of lung sections collected 21 days after bleomycin injection (11 days after AAV6.2FF-miR-124 administration) are shown. (F–I) ATII cells were stained for pro-SPC (white), CLDN4 (red), and GFP (green), with nuclei counterstained with Hoechst (blue). Quantitative image analysis includes the percentage of GFP⁺ cells over total cells (G), SPC⁺/GFP⁻ cells over total cells (H), and GFP⁺/CLDN4⁺/SPC⁺ triple-positive cells over total cells (I). (J–M) ATI cells were stained for podoplanin (white), GFP (green), and CLDN4 (red), with nuclei counterstained with Hoechst (blue) to visualize lineage-labeled ATI cells. Quantification includes GFP⁺/podoplanin⁺ cells over total cells (K), GFP⁺/CLDN4⁺/podoplanin⁺ cells over total cells (L), and GFP⁻/CLDN4⁺/podoplanin⁺ cells over total cells (M). Statistical significance for multiple comparisons was determined using one-way ANOVA followed by Dunnett’s multiple comparisons test, whereas pairwise comparisons were analyzed using an unpaired two-tailed Welch’s t-test.. *P < 0.05; **P < 0.01; ***P < 0.0001.

We next asked whether the ATII cell isolation protocol also enabled the recovery of ATII progenitor cells or immature ATII cell populations. ATII cells were isolated using a negative selection–based protocol involving antibody-conjugated magnetic beads, size-exclusion filtering, and pre-plating, as described in the Methods section. We characterized this ATII-enriched population, isolated from tamoxifen treated SFTPC-CreER[T2]/mTmG mice, by flow cytometry and found that among EPCAM+ cells, approximately 80.7% were GFP^+^, while 15.2% were MHC-II+/GFP-negative (Supplementary Fig. 7A). EPCAM+MHC-II+ cells were previously identified by Strunz et al. ^8^ as a subpopulation of club cells termed MHC-II+ club cells. This subpopulation has been shown to support alveolar repair by differentiating into mature ATII and ATI cells ^12^. Therefore, we conclude that our ATII isolation protocol includes approximately 15% MHC-II+/GFP-negative cells, which upon treatment with hsa-miR-124-3p, could potentially give rise to the observed RAGE+/GFP-negative cells.

### AAV6.2FF targets ATII cells, progenitors, and injury-induced epithelial intermediates *in vivo*

Given the therapeutical efficacy of AAV6.2FF-miR-124 in the bleomycin mouse model of lung fibrosis, and the *in vitro* results supporting effect of miR-124-3p on ATII progenitors, we aimed to demonstrate that AAV6.2FF can target *in vivo* the relevant cell types that support effective alveolar repair, beyond SPC⁺ ATII cells. In particular, we focused on progenitors MHC-II⁺ CLUB cells and KRT8⁺/SPC⁻ aberrant intermediates.

We therefore intratracheally injected C57BL/6 mice with AAV6.2FF-GFP. Ten days after injection, mice were sacrificed and lungs were processed for histological analysis (Experimental Scheme in Supplementary Fig. 8A). We confirmed that AAV6.2FF-GFP can target 6,7% ± 1.1% of MHC-II⁺/SPC⁻ cells (Supplementary Fig. 8B,C) quantified by IF staining and 14.6% ± 7,4% by cytofluorimetry (Supplementary Fig.8G). Similarly, we injected AAV6.2FF-GFP into C57BL/6 mice ten days after bleomycin administration (Experimental Scheme in Supplementary Fig. 8D). Lungs collected ten days post-injection showed that 21.7% ± 4.5% of KRT8⁺/SPC⁻ cells were GFP-positive by immunofluorescence staining (Supplementary Fig. 8E,F) and by cytofluorimetry 33.2% ± 4.8% (Supplementary Fig. 8H).

Finally, using tamoxifen-treated SFTPC-CreER[T2]/mTmG mice, we demonstrated that intratracheal delivery of AAV6.2FF-miR-124 effectively induced overexpression of hsa-miR-124-3p in MHC-II⁺ Club distal progenitor cells (Suppl. Fig. 9A,B). These findings further confirm that AAV6.2FF-mediated gene transfer can effectively target the relevant cell populations that support epithelial-driven alveolar repair.

### AAV6.2FF-miR-124 reverses bleomycin-induced lung fibrosis by promoting epithelial-driven alveolar repair

As shown in the AAV6.2FF-miR-124 efficacy study (Fig. 3), overexpression of hsa-miR-124-3p in alveolar epithelial type II (ATII) cells is sufficient to reverse disease progression, resulting in evident preservation of lung architecture. However, it remained unclear whether this effect was driven by enhanced ATII-mediated alveolar repair. To address this, we replicated the therapeutic study from Fig. 3 using tamoxifen-treated SFTPC-CreER[T2]/mTmG mice, which allowed us to track ATII-to-ATI cell transdifferentiation upon treatment and over time. At 11 days post-AAV administration (21 days post-bleomycin), we observed that, consistent with our *in vitro* findings, miR-124-3p primarily targets pre-ATII progenitors that were GFP⁻ at the time of administration. This resulted in a modest, non-significant increase in the total number of GFP⁺/SPC⁺ cells (Fig. 4G,F), accompanied by a significant increase in SPC^+^/GFP⁻ cells (Fig. 4I,F). These findings suggest that miR-124-3p stimulates alveolar repair at the level of ATII progenitors, enhancing their differentiation toward ATII cells. By probing this process at a single time point, we detected an increased number of newly formed SPC⁻ cells indicative of active transdifferentiation.

Interestingly, we confirmed that the increase in SPC⁺/CLDN4⁺ intermediate cells observed in C57BL/6 mice (Fig. 3I) is mainly sustained by mature SPC⁺ cells, as demonstrated by the increase in SPC⁺/GFP⁺/CLDN4⁺ cells (Fig. 4F,I). In line with this, we observed a marked increase in PDPN⁺/GFP⁺ cells (Fig. 3J,K), suggesting that miR-124-3p act also on mature ATII cells. Of note, we also detected an increase in PDPN⁺/CLDN4⁺ cells, originating from both GFP⁺ ATII-derived and GFP⁻ lineages, indicating effective formation of tight junctions and restoration of alveolar barrier integrity, regardless of the cellular origin of the newly formed ATI cells (Fig. 4J,L,M).

In conclusion, these findings support a model in which hsa-miR-124-3p acts at multiple levels, targeting both ATII progenitors and mature ATII cells to stimulate alveolar repair. Importantly, by engaging progenitor populations, miR-124-3p may help replenish and maintain the ATII pool, thereby preventing its depletion during sustained injury. This coordinated action enables continued differentiation toward ATI cells without exhausting the ATII compartment, ultimately supporting the restoration of an intact alveolar epithelial barrier.

### Endogenous miR-124-3p drives stage-specific regulation of ATII cell fate and plasticity through EZH2 and C/EBP**α** targeting

To better understand the mechanism of action of miR-124-3p, we first investigated whether miR-124-3p could physiologically contribute to alveolar repair following injury in the adult lung. To address this, we exploited the spontaneous resolution phase of the bleomycin mouse model of lung fibrosis, which occurs between 45 and 60 days after injury ^50^. As shown in Fig. 5A, endogenous miR-124-3p expression was analyzed at multiple time points following bleomycin-induced lung injury and remodelling. miR-124-3p levels increased after injury, peaked at day 21, and declined during fibrosis resolution, consistent with the pattern observed by Masson’s trichrome staining (Fig. 5B,C). Consistently, aged 20-month-old bleomycin-treated mice, which are known to display defective alveolar repair ^36^, failed to upregulate miR-124-3p at day 21 compared with young mice (Fig. 5D-F).

**Figure 5.**
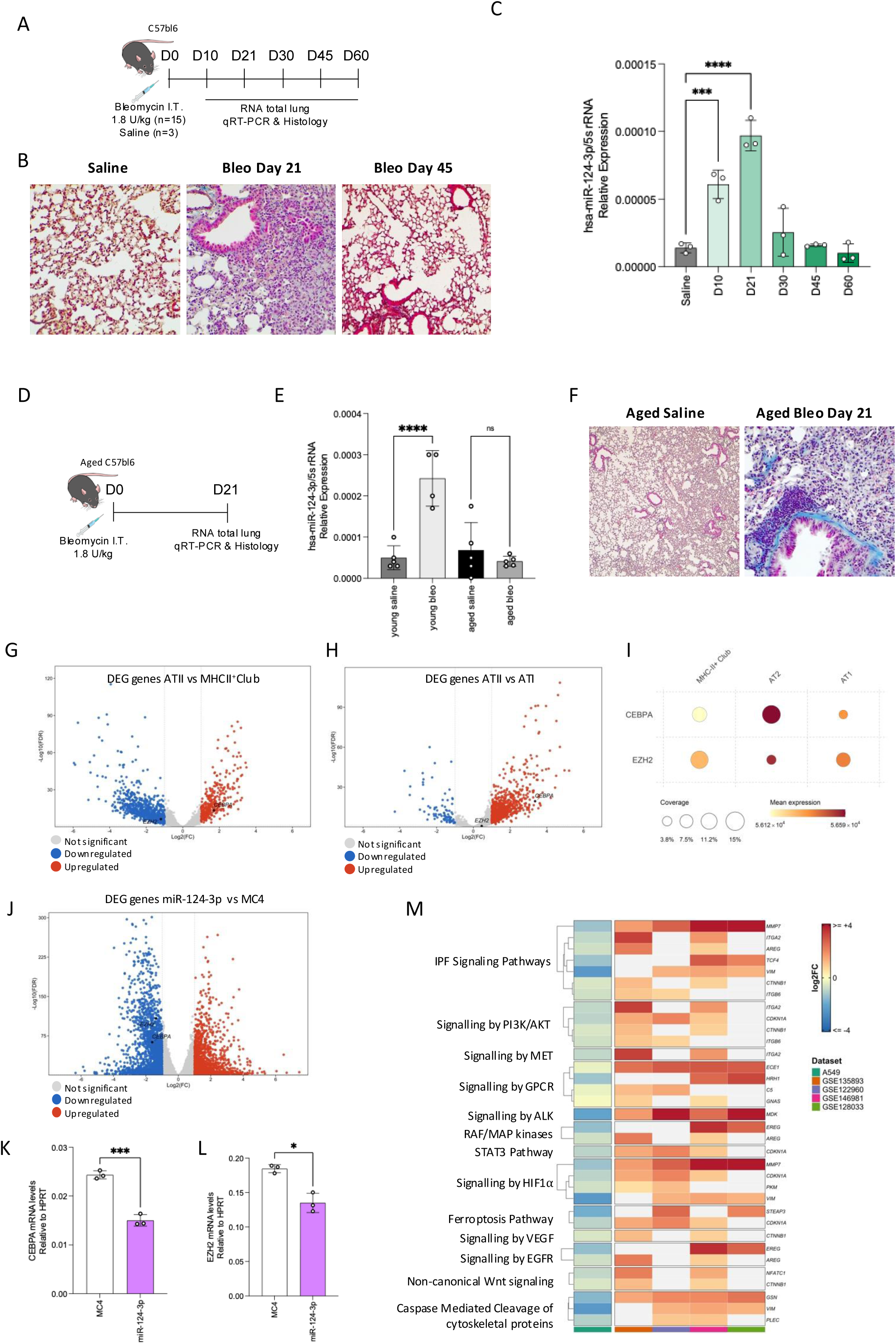
Endogenous miR-124-3p is associated with stage-specific regulation of ATII cell fate and plasticity through EZH2 and C/EBPα targeting. (A) Schematic overview of the experimental design to evaluate miR-124-3p expression across the different phases of the young bleomycin-induced mouse model of IPF. Mice were sacrificed 10 days after bleomycin injection (inflammatory phase), 21 days (peak fibrosis), and 30, 45, and 60 days (damage resolution phases). (B) Representative Masson’s trichrome staining of lung sections from saline-treated mice and from mice sacrificed 21 and 45 days after bleomycin injection. (C) miRCURY LNA miRNA–based quantitative PCR analysis of mature miR-124-3p expression in total lung tissue at 10, 21, 30, 45, and 60 days after bleomycin administration. (D) Schematic overview of the experimental design to compare miR-124-3p expression levels in aged (20-month-old) versus young mice treated with bleomycin. All mice were sacrificed 21 days after bleomycin injection. Lungs were harvested; the right lobe was processed for RNA extraction, whereas the left lobe was used for histological analysis (Masson’s trichrome staining). (E) miRCURY LNA miRNA–based quantitative PCR analysis of mature miR-124-3p expression in total lung tissue from saline-treated young and aged mice, and from mice 21 days after bleomycin injection. (F) Representative Masson’s trichrome staining of lung sections from aged mice treated as in panel D. (G–H) Volcano plots of differentially expressed genes (DEGs) comparing ATII vs MHC-II⁺ Club cells (G) and ATII vs ATI cells (H), from published mouse scRNAseq study (GSE141259). Up- and down-regulated genes are colored in red and blue, respectively. C/EBPα and EZH2 are highlighted in bold. (I) Dot plot showing the expression of C/EBPα and EZH2 across three alveolar epithelial populations (MHC-II⁺ Club, ATII, and ATI cells) from the dataset described in (G–H). Dot size represents the fraction of cells expressing the gene within each population, and color intensity encodes the scaled average expression level (z-score of log-normalized counts). (J) Volcano plot of differentially expressed genes (DEGs) in A549 transfected with miR-124-3p mimic vs mimic control (MC4). Up- and down-regulated genes are colored in red and blue, respectively. C/EBPα and EZH2 are highlighted in bold. (K–L) RT-qPCR quantification of C/EBPα (K) and EZH2 (L) mRNA levels in A549 cells transfected with miR-124-3p mimic or mimic control (MC4). Total RNA was extracted 72 h post-transfection, reverse-transcribed, and expression was normalized to HPRT using the 2^(–ΔΔCt) method. Each point represents a different biological replicates. (M) Heatmap of genes that are simultaneously (i) down-regulated in miR-124-3p–transfected A549 cells relative to MC4 control and (ii) up-regulated in ATII cells of IPF versus healthy donor lungs across published human scRNAseq datasets (GSE135893, GSE12296, GSE146981, GSE12033). Overlapping genes were further filtered to retain those belonging to the top canonical pathways predicted to be inhibited by Ingenuity Pathway Analysis (IPA, QIAGEN) on the miR-124-3p A549 DEG signature. Each row corresponds to a single gene. Genes have been clustered by pathway. Statistical significance was determined using one-way ANOVA. For comparisons against a single control group, Dunnett’s multiple comparisons test was applied. For selected pairwise comparisons between specific groups, Šídák’s multiple comparisons test was used. *P < 0.05; **P < 0.01; ***P < 0.0001.

These findings support a role for miR-124-3p during physiological alveolar regeneration following bleomycin-induced lung injury. Because alveolar regeneration recapitulates key features of developmental alveologenesis ^38,51^, we next considered whether miR-124-3p might participate in regulatory pathways governing distal epithelial cell fate. Previous studies have reported dynamic miR-124-3p expression during fetal lung development^37^ and identified a developmental trajectory linking SOX9+ progenitors, nascent AT2 cells, and mature AT2 cells through coordinated changes in EZH2 and C/EBPα expression ^39^. Furthermore, prior evidence supports that miR-124-3p directly targets both *EZH2* and C/EBPα in distinct biological contexts ^52,53^.

Our data are consistent with a similar regulatory mechanism in lung epithelial cells: overexpression of a miR-124-3p mimic in A549 cells resulted in significant downregulation of both EZH2 and C/EBPα at the mRNA level (Fig. 5J–L). In this context, reanalysis of the single-cell dataset from Strunz *et al.* ^8^ revealed expression patterns of EZH2 and C/EBPα in MHC-II⁺ Club distal progenitors and AT2 cells fully concordant with those described by Sawhney *et al.* ^39^: EZH2 expression was markedly lower in ATII cells relative to MHC-II⁺ Club distal progenitors (Fig. 5G,I), whereas C/EBPα followed the inverse gradient, being higher in AT2 vs both MHC-II⁺ Club distal progenitors (Fig.5G,I) and ATI cells (Fig. 5H,I). Moreover, the efficient in vivo transduction of both MHC-II⁺ Club distal progenitor cells and SPC⁺ AT2 cells by AAV6.2FF-miR-124 (Supplementary Figs. 6, 9) suggests that the therapeutic effects observed following miR-124-3p delivery *in vivo* may be mediated, at least in part, through modulation of the above described EZH2 - C/EBPα-regulated distal epithelial differentiation axis.

Last, to assess the broader translational relevance of miR-124-3p-mediated transcriptional remodelling in human alveolar epithelial cells, we intersected all genes downregulated upon hsa-miR-124-3p overexpression in A549 cells with genes consistently upregulated in IPF patients across at least two out of four publicly available human single-cell RNA-seq datasets, including Habermann/GSE135893 ^7^, Reyfman/GSE122960 ^54^, Yao/GSE146981 ^55^, and Morse/GSE128033 ^56^ (Supplementary Fig. 10A). This analysis identified 173 genes, the majority of which were consistently upregulated across the analyzed IPF datasets and therefore inversely regulated by miR-124-3p overexpression. In contrast, a smaller subset of 55 genes was concordantly downregulated across all datasets; however, pathway enrichment analysis indicated that these genes were not associated with established IPF-related biological processes (Supplementary Fig. 10C).

To further contextualize these findings, we interrogated this gene set against canonical pathways previously associated with IPF pathogenesis using Ingenuity Pathway Analysis (IPA) (Supplementary Fig. 10B) and Reactome databases, additionally incorporating two recently described epithelial remodelling and transitional cell state pathways (R-HSA-264870 and R-HSA-372790) ^57^ ^58^. This approach identified a final subset of 34 genes linked to multiple fibrosis-associated pathways, all of which were upregulated in at least two independent human IPF single-cell datasets (Fig. 5M).

Taken together, these findings support a role for miR-124-3p in distal epithelial cell-state transitions associated with alveolar repair. Beyond regulation of the EZH2– C/EBPα axis, integration of transcriptomic and human single-cell datasets suggests that miR-124-3p influences multiple epithelial gene programs and fibrosis-associated pathways dysregulated in IPF, highlighting its potential as a pleiotropic regulator of lung epithelial repair.

## Discussion

The treatment of IPF remains a major unmet clinical need. Lung transplantation is currently the only intervention capable of replacing lost lung tissue with functional alveoli, but it is feasible only for a minority of patients. Current pharmacological treatments can slow functional decline; however, they cannot replace the lost alveolar surface or restore functional alveolar architecture. These limitations highlight the need for regenerative strategies that promote endogenous alveolar repair, reflecting a broader paradigm shift in fibrosis research from suppression of fibrogenesis toward resolution and tissue restoration^59^.

Over the past decade, accumulating evidence has established epithelial dysfunction, rather than primary fibroblast activation, as a central driver of idiopathic pulmonary fibrosis (IPF), marking a fundamental shift in disease pathogenesis ^15^. In particular, impaired differentiation of alveolar epithelial type II (ATII) cells into alveolar epithelial type I (ATI) cells has emerged as a key pathological feature that sustains aberrant epithelial states and promotes fibrotic remodelling ^2,7,8^.

Despite this paradigm shift, approved and emerging pharmacological strategies remain largely antifibrotic rather than pro-regenerative. Pirfenidone, nintedanib, and the recently developed PDE4B inhibitor Nerandomilast mainly act by modulating profibrotic, inflammatory, vascular, and tissue-remodelling pathways. These approaches are clinically important, but they are expected primarily to slow disease progression rather than rebuild the damaged alveolar epithelium or restore lung architecture ^60,61^.

In this study, we conducted the first unbiased, large-scale functional microRNA screen specifically designed to interrogate ATII-to-ATI transdifferentiation, using 2,042 human microRNA mimics in primary mouse ATII cells. By directly assessing epithelial fate outcomes rather than expression changes, this approach enabled the identification of microRNAs with functional relevance to alveolar regeneration. By directly measuring epithelial fate outcomes rather than relying only on transcriptomic changes, this approach identified miR-124-3p as the most effective and biologically compelling candidate in promoting ATII-to-ATI transdifferentiation in vitro. Subsequent validation showed that miR-124-3p enhanced ATI-like differentiation in bleomycin-exposed ATII cells, while showing no pro-fibrotic effect on TGF-β-stimulated lung fibroblasts and no proliferative effect in A549 cells.

The in vivo data support the therapeutic relevance of this screening strategy. Using the engineered AAV6.2FF capsid ^33,34^, we provided direct evidence that epithelial-specific modulation of microRNA signaling is sufficient to promote lung repair without directly targeting fibroblasts in the bleomycin-induced lung fibrosis model. Notably, AAV6.2FF efficiently transduced not only mature ATII cells but also MHC-II⁺ club distal progenitor cells and injury-induced KRT8⁺ epithelial intermediates, enabling coordinated modulation of epithelial programs across multiple stages of the regenerative hierarchy, an advantage in the context of fibrotic lung disease. Consistent with these findings, *in vivo* lineage-tracing experiments support a direct contribution of mature ATII cells and epithelial progenitors to miR-124-3p– driven alveolar repair. These effects occurred without deliberate targeting of fibroblasts, supporting the concept that restoration of epithelial regenerative programmes can indirectly reshape the fibrotic niche.

Such pleiotropic and coordinated actions are well aligned with the observation that endogenous miR-124-3p expression dynamically increases following bleomycin-induced injury further supporting the possibility that miR-124-3p participates in physiological alveolar repair programs. Interestingly, aged mice, which display defective alveolar regeneration ^36^, failed to upregulate miR-124-3p after bleomycin administration. This finding is consistent with the broader concept outlined above that age-associated failure to activate epithelial repair programmes contributes to defective regeneration in fibrotic lung disease.

These results are particularly intriguing in light of the established parallels between alveolar regeneration and developmental alveologenesis. ^38,51^. Wang et al. previously showed that miR-124 is highly expressed during early fetal lung and peaks at E16 to then progressively declines during epithelial maturation ^37^. Interestingly, this developmental stage closely corresponds to the emergence of nascent AT2 (nAT2) cells described by Sawhney et al. ^39^, which are characterized by a highly plastic cellular state associated with high EZH2 and low C/EBPα expression. As development proceeds, progressive transition toward the mature AT2 (mAT2) state and loss of epithelial plasticity corresponds to induction of C/EBPα. Importantly, Sawhney et al. further demonstrated that transient C/EBPα downregulation is required in the adult lung to reactivate AT2 plasticity and enable ATII-to-ATI transdifferentiation during alveolar repair. Accordingly, Lee et al. have also implicated EZH2 in aberrant epithelial remodelling and maladaptive repair responses following adult lung injury.^40^.

In this context, our findings are consistent with a model in which miR-124-3p exerts context-dependent functions across distinct epithelial cell states. In progenitor-like epithelial populations high for EZH2 expression, miR-124-3p may facilitate epithelial maturation programs by targeting EZH2 and enabling C/EBPα transcriptional programs, whereas in mature ATII cells it may promote reparative plasticity through transient modulation of C/EBPα-dependent transcriptional programs. Consistently, reanalysis of the Strunz et al ^8^ single-cell dataset revealed expression patterns of EZH2 and C/EBPα in MHCII-Club distal progenitors and ATII cells that closely mirrored those described by Sawhney et al. ^39^, with higher EZH2 expression in progenitor-like epithelial populations and higher C/EBPα expression in mature ATII cells. Together, these observations suggest that miR-124-3p does not enforce a single epithelial state fate, but rather coordinates distinct regenerative programs across multiple stages of the alveolar epithelial hierarchy.

The transcriptomic integration with multiple public human IPF single-cell RNA-seq datasets further suggested that miR-124-3p broadly alleviates pathological pressure on epithelial cells by downregulating multiple genes associated with fibrosis-related pathways in human IPF. Although A549 cells cannot substitute for primary human ATII cells, these data support the idea that miR-124-3p may act as a pleiotropic regulator of epithelial programmes relevant to human fibrotic remodelling, beyond the EZH2-C/EBPα axis alone.

This study has several limitations. First, the functional screen was performed in isolated 2D-cultured ATII cells and therefore does not fully capture the multicellular, biomechanical, and inflammatory context of the injured alveolar niche. Future discovery efforts would benefit from in vivo functional screening approaches in three-dimensional alveolar organoids or lung on chip devices, possibly employing the use of IPS-derived human cells ^62^.

Second, in vivo validation relied on viral-mediated microRNA overexpression. While the use of the AAV6.2FF capsid enabled preferential targeting of alveolar epithelial cells, providing strong evidence that epithelial-specific modulation of microRNA activity is sufficient to attenuate fibrotic remodelling without directly targeting fibroblasts, non-viral delivery strategies will ultimately be required to improve translational feasibility.

Third, miR-124-3p delivery represents a gain-of-function approach; future unbiased functional screening studies aimed at identifying pathogenic microRNAs negatively regulating ATII-to-ATI transdifferentiation may enable loss-of-function therapeutic strategies with potentially improved safety and translatability.

Last, the present study primarily relied on histological, biochemical, and lineage-tracing endpoints. Future studies should assess whether miR-124-3p-mediated epithelial repair translates into measurable improvements in lung mechanics and gas-exchange function.

Together, these findings establish miR-124-3p as a previously unrecognized endogenous regulator of alveolar epithelial cell plasticity in the adult mouse lung, capable of driving alveolar repair in the setting of established pulmonary fibrosis. More broadly, this work highlights the value of unbiased, function-driven microRNA screening in identifying epithelial regulators that may enable the development of pro-regenerative therapies designed to complement existing antifibrotic treatments.

## Materials and Methods

### ATII cell isolation and culture

Alveolar type II (ATII) epithelial cells were isolated from adult (8-week-old) C57BL/6 mice. Lungs were perfused with PBS, after which 1 mL of Dispase (SIAL-Corning, #354235) was instilled intratracheally. The lungs, were then immersed in an additional 2 mL Dispase solution and incubated for 30 min at room temperature. Distal lung tissue was mechanically dissociated using a McIlwain tissue chopper (Metrohm, USA) and incubated with DNase I (20 µg mL⁻¹; Sigma-Aldrich #04536282001) for 10 min at 37 °C. The resulting cell suspension was sequentially filtered through 100 µm (ClearLine cell strainer #141380C) and 40 µm nylon strainers (ClearLine cell strainer #141378C). The cell pellet was resuspended in Dulbecco’s modified Eagle’s medium (DMEM, Gibco, #21885108) with GlutaMAX™ I, supplemented with 10% fetal bovine serum (FBS; Euroclone # ECS5000L) and antibiotics (Pennicillin-Streptomycin, Thermo Fisher Scientific, #15140-122), and subjected to two rounds of differential adherence on plastic dishes (1 h each) to deplete fibroblasts and macrophages. Immune cells were removed by negative magnetic selection using the Dynabeads™ Untouched™ Mouse T Cell Kit (Thermo Fisher Scientific, #11413D) and the Dynabeads™ Mouse DC Enrichment Kit (Thermo Fisher Scientific, #11429D), following the manufacturer’s instructions. Purified ATII cells were seeded onto plates pre-coated with fibronectin (1 mg mL⁻¹; Life Technologies, #33010018) and bovine gelatin (0.2%; Sigma-Aldrich, G9391), and were maintained in PneumaCult™-ALI Medium(Stemcell Technologies, #05050) supplemented with 10% FBS and antibiotics under standard culture conditions (37 °C, 5% CO₂).

### Primary lung fibroblast isolation and culture

Primary lung fibroblasts were isolated from adult *C57BL/6* mice (8 weeks old). Lungs were perfused via the right ventricle with sterile PBS to remove blood, then mechanically minced into small fragments using sterile scissors. Tissue pieces were digested using the Skeletal Muscle Dissociation Kit (Miltenyi Biotec, #130-098-305) according to the manufacturer’s protocol. Briefly, fragments were incubated for 30 min at 37 °C under gentle agitation in an enzyme mix consisting (per gram of tissue) of 200 µL Enzyme D, 50 µL Enzyme P, and 36 µL Enzyme A. The digested material was passed through a 70 µm cell strainer (ClearLine cell strainer #141379C) and centrifuged to collect the cell pellet. Cells were resuspended in DMEM with GlutaMAX™ I (Gibco, #218851), supplemented with 10% FBS and 1% penicillin-streptomycin-amphotericin B, and incubated at 37 °C, 5% CO₂. After 4 h, the medium was replaced to remove non-adherent cells. Fibroblasts were reverse-transfected 48 h after seeding.

### A549 Cell culture

A549 human alveolar epithelial cells were cultured in DMEM supplemented with GlutaMAX™ I (Gibco, #218851), sodium pyruvate, pyridoxine, and 1 g L⁻¹ glucose, together with 10% FBS (Euroclone, # ECS5000L) and 1% penicillin streptomycin (Thermo Fisher Scientific, #15140-12). Cells were routinely passaged at 70–80% confluence under standard conditions (37 °C, 5% CO₂).

### High-throughput microRNA screening

Due to the limited number of primary ATII cells that could be obtained from a single isolation experiment, the screen was performed in seven independent experiments, each corresponding to one of the seven library plates comprising the complete miRNA screening library. Primary ATII cells were reverse-transfected with an arrayed library containing 2042 human miRNA mimics at a final concentration of 25 nM. As a negative control, cells were transfected with cel-miR-231-3p (non-targeting control, MC4), while siUBC was included as a positive control to monitor transfection efficiency. Prior to transfection, 384-well PhenoPlates were coated with fibronectin (1 mg mL⁻¹; Thermo Fisher Scientific, #33010018) and bovine gelatin (0.2%; Sigma-Aldrich, #G9391). Following coating, the miRNA library was arrayed onto the plates using a Hamilton STARlet liquid handler, dispensing 5 µL of 500 nM miRNA mimic solution into each well. Subsequently, a transfection mixture consisting of 15 µL Opti-MEM and 0.15 µL Lipofectamine RNAiMAX was added to each well using a MultiDrop Combi reagent dispenser (Thermo Fisher Scientific). After a 30-minute incubation at room temperature to allow complex formation, 15,000 cells per well, suspended in 50 µL PneumaCult™-ALI Medium (Stemcell Technologies, #05050) t) supplemented with 10% fetal bovine serum (Euroclone, # ECS500) and antibiotics, were dispensed into each well. Cells were then cultured under standard conditions (37°C, 5% CO₂) for 72 hours. At the end of the incubation period, plates were washed with PBS and fixed with 4% paraformaldehyde (PFA, Societa’ Italiana Chimici #15714-S) for 10 minutes at room temperature. Cells were subsequently subjected to immunofluorescence staining for RAGE and p21 using the antibodies listed below and following the protocol described in the dedicated Immunofluorescence Staining section. Plates were imaged using an Operetta CLS high-content imaging system (Revvity) equipped with a 20× objective (NA 0.4). A total of nine fields of view per well were acquired. Image analysis was performed using Signals Image Artist software (Revvity). The following parameters were quantified for each well: total cell number, number of RAGE-positive cells, and number of p21-positive cells. For each plate, absolute cell counts were normalized to the corresponding cel-miR-231-3p negative control wells. Normalized values were then used to calculate Z-scores for each measured parameter, enabling comparison across the screening dataset. All primary hits were subsequently subjected to manual image review to exclude candidates arising from imaging or staining artifacts. To further account for potential inter-experimental variability associated with independent primary ATII cell isolations, the top-performing microRNA candidates from each screening plate were selected for downstream validation and characterization.

### Transfection in 96-well plates

microRNA mimics were spotted into 96-well plates and pre-incubated with Opti-MEM and Lipofectamine RNAiMAX (Thermo Fisher Scientific, #13778030), according to manufacturer’s instructions, at room temperature to allow transfection complex formation. Primary ATII cells (50,000 cells/well) and A549 cells (2,500 cells/well) were then seeded onto the complexes reaching a final microRNA concentration of 25 nM, while primary murine lung fibroblasts (5,000 cells/well) were transfected reaching a final concentration of 12.5 nM. The final assay volume per well was 150 µL (50 µL transfection complex + 100 µL cell suspension). Cells were cultured under standard conditions (37°C, 5% CO₂) for 72 hours.

### In vitro Proliferation assay

A549 cells and primary murine fibroblasts were seeded in 96-well plates and transfected as described above. Four hours before fixation, the medium was replaced with fresh medium containing 10 mM 5-ethynyl-2′-deoxyuridine (EdU) to label proliferating cells. Cells were fixed in 4% PFA for 20 min and stained with primary antibodies, DAPI, and EdU using the Click-iT™ EdU Cell Proliferation Kit for Imaging (Thermo Fisher Scientific, #C10338) according to the manufacturer’s recommendations.

### In vitro TGF-β treatment of primary murine lung fibroblasts

Two days after isolation, primary murine lung fibroblasts cultured in high-glucose DMEM with 10% FBS were detached and reseeded (5 × 10³ cells per well) in DMEM containing 0.5% FBS. After 24 h, cells were treated with TGF-β1 (2 ng mL⁻¹; PeproTech, #100-21C-10UG, Thermo Fisher Scientific) for 72h.

### Immunofluorescence staining on fixed cells

At the desired time point, cells were fixed in 4% PFA (Societa’ Italiana Chimici #15714), permeabilized with 0.1% Triton X-100 (Thermo Fisher Scientific, #A16046.AE) for 15 min, and blocked in 5% BSA (Merck Life Science #A2153) added of 0.1% Triton X-100 for 1 h at room temperature. And them incubated with the selected primary antibodies (Table 1) O/N at 4°C. Following three washes with PBS, cells were incubated with the appropriate fluorophore-conjugated secondary antibodies (Table 2) for 2 h at RT, protected from light. Nuclei were counterstained with Hoechst 33342 (Thermo Fisher Scientific # H3570) (1:5000) for 10 min at RT.

**Table 1.**
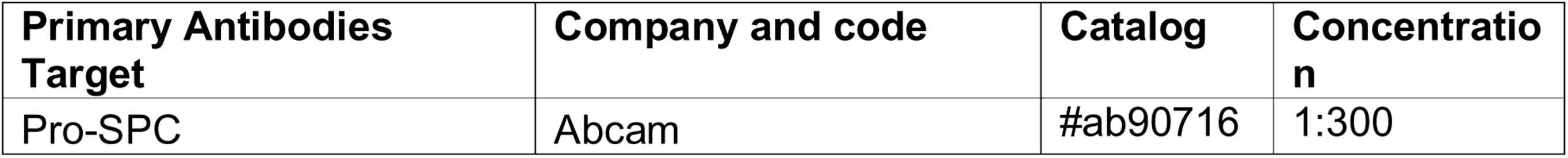
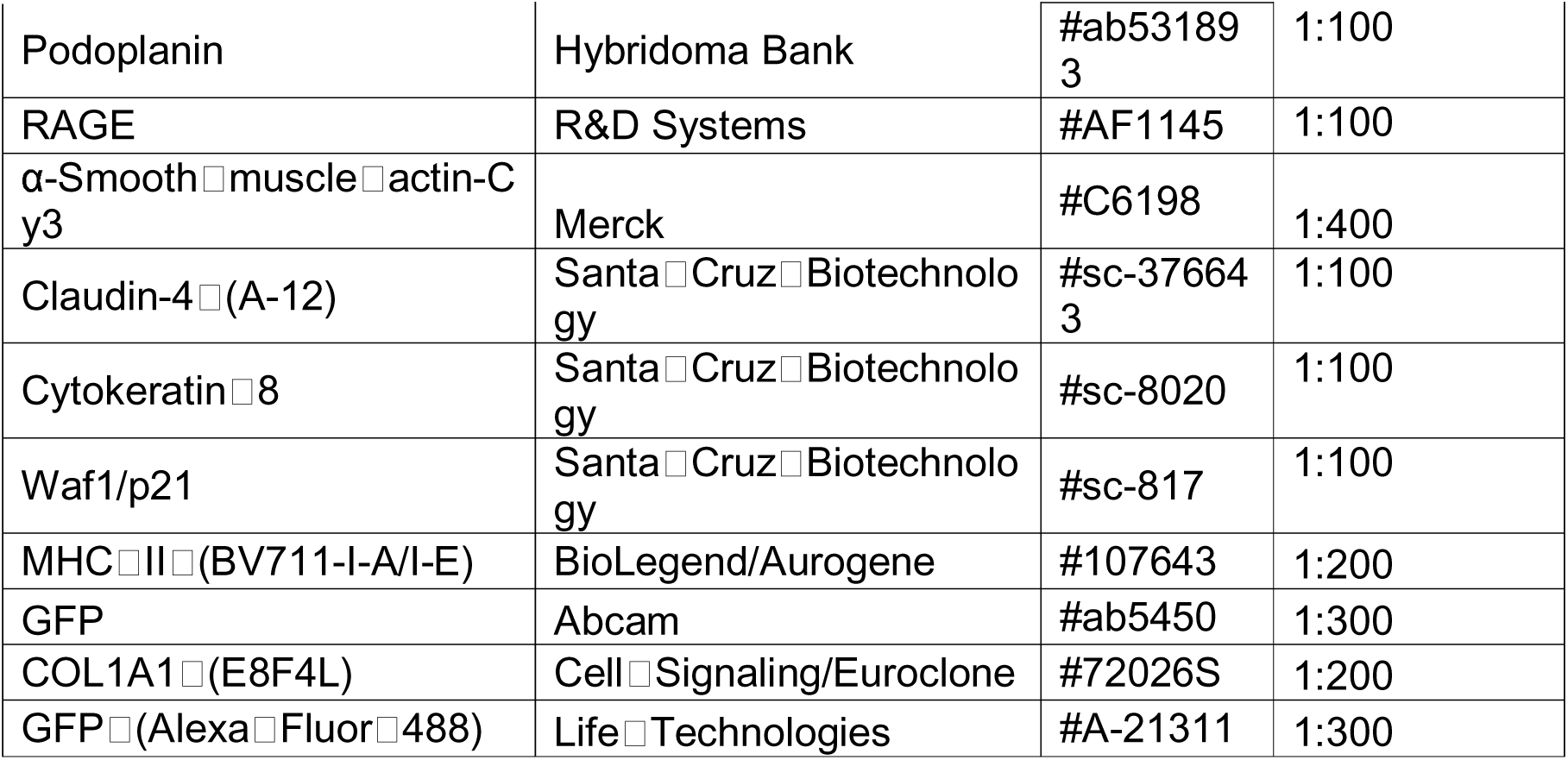
Primary Antibodies information and working concentration.

| Primary Antibodies<br>Target | Company and code | Catalog | Concentratio<br>n |
| --- | --- | --- | --- |
| Pro-SPC | Abcam | #ab90716 | 1:300 |
| Podoplanin | Hybridoma Bank | #ab531893 | 1:100 |
| RAGE | R&D Systems | #AF1145 | 1:100 |
| α-Smooth muscle actin-Cy3 | Merck | #C6198 | 1:400 |
| Claudin-4 (A-12) | Santa Cruz Biotechnology | #sc-376643 | 1:100 |
| Cytokeratin 8 | Santa Cruz Biotechnology | #sc-8020 | 1:100 |
| Waf1/p21 | Santa Cruz Biotechnology | #sc-817 | 1:100 |
| MHC II (BV711-I-A/I-E) | BioLegend/Aurogene | #107643 | 1:200 |
| GFP | Abcam | #ab5450 | 1:300 |
| COL1A1 (E8F4L) | Cell Signaling/Euroclone | #72026S | 1:200 |
| GFP (Alexa Fluor 488) | Life Technologies | #A-21311 | 1:300 |

**Table 2.** Secondary Antibodies information.

| Secondary Antibodies (target) | Company and code | Catalog | Concentration |
| --- | --- | --- | --- |
| Alexa Fluor™ 568 Donkey anti goat IgG | Invitrogen | #A-11057 | 1:500 |
| Alexa Fluor™ 488 Donkey anti goat IgG | Invitrogen | #A-11057 | 1:500 |
| Alexa Fluor™ 647 Donkey anti goat IgG | Invitrogen | #A-21447 | 1:500 |
| Alexa Fluor™ 568 Donkey anti mouse IgG | Invitrogen | #A-10037 | 1:500 |
| Alexa Fluor™ 488 Donkey anti mouse IgG | Invitrogen | #A-212062 | 1:500 |
| Alexa Fluor™ 647 Donkey anti mouse IgG | Invitrogen | #A-31571 | 1:500 |
| Alexa Fluor™ 594 Donkey anti rabbit IgG | Invitrogen | #A-21207 | 1:500 |
| Alexa Fluor™ 488 Donkey anti rabbit IgG | Invitrogen | #A-21206 | 1:500 |
| Alexa Fluor™ 647 Donkey anti rabbit IgG | Invitrogen | #A-31576 | 1:500 |
| Goat Anti-Syrian Hamster IgG H&L (Alexa Fluor 488) | Abcam | #ab180063 | 1:500 |
| Goat Anti-Syrian Hamster IgG H&L (Alexa Fluor 647) | Abcam | #ab1801117 | 1:500 |

### Immunofluorescence staining on Tissue Slides

Lungs were fixed overnight in 4% PFA at 4 °C, transferred to 70% ethanol, paraffin-embedded, and sectioned (5 µm). Sections were deparaffinized at 60 °C for 2 h, incubated in xylene (30 min) (Bio-optica, #26-100105), rehydrated through ethanol gradients (100%, 90%, 75%, 50%), and washed. Antigen retrieval was performed in 10 mM sodium citrate buffer (0.05% Tween-20, pH 6) at 100 °C for 30 min, followed by permeabilization in 0.1% Triton X-100/PBS (20 min, RT). Slides were blocked in 5% BSA/0.1% Triton-PBS (1 h, RT), incubated overnight at 4 °C with primary antibodies (Table 1), washed, then incubated with fluorescent secondary antibodies (Table 2) for 2 h at RT in the dark. Slides were counterstained with Hoechst 33342 (1:5000, 20 min) and mounted in Mowiol 40-88 (Merck Life Science, #324590).

### Quantitative real-time PCR (qRT-PCR)

Total RNA was extracted using the miRNeasy Micro Kit or miRNeasy Mini Kit (Qiagen, #217084 and #217004, respectively), according to the manufacturer’s instructions. cDNA was synthesized using the M-MLV Reverse Transcriptase Kit (Thermo Fisher Scientific, #28025-013), and quantitative real-time PCR (qRT-PCR) was performed on a C1000 Thermal Cycler (Bio-Rad). Relative gene expression was normalized to the reference genes Tuba, 18S rRNA, and Hprt, as appropriate. Primer sequences used for qRT-PCR amplification are listed in Table 3.

**Table 3.**
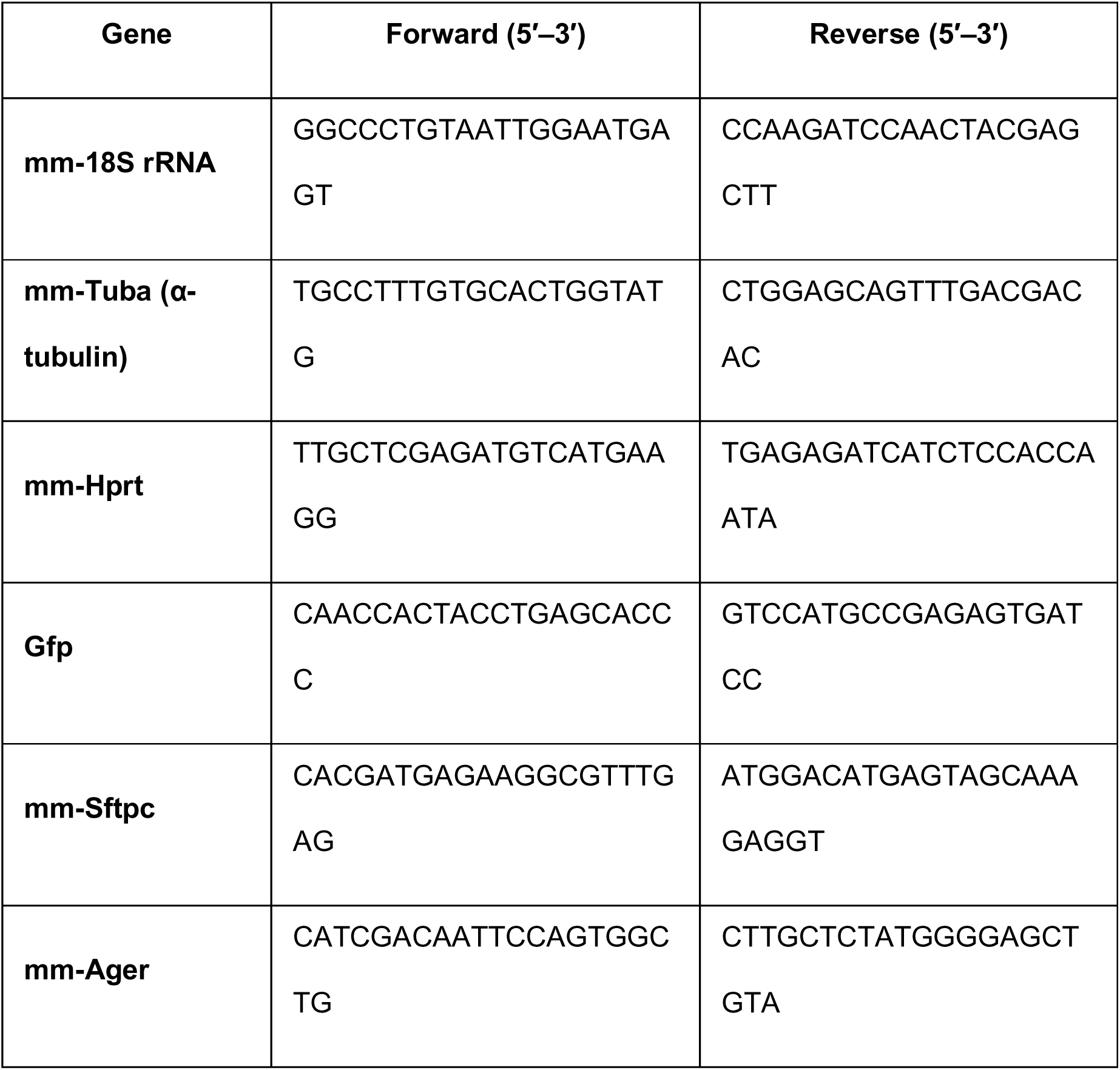
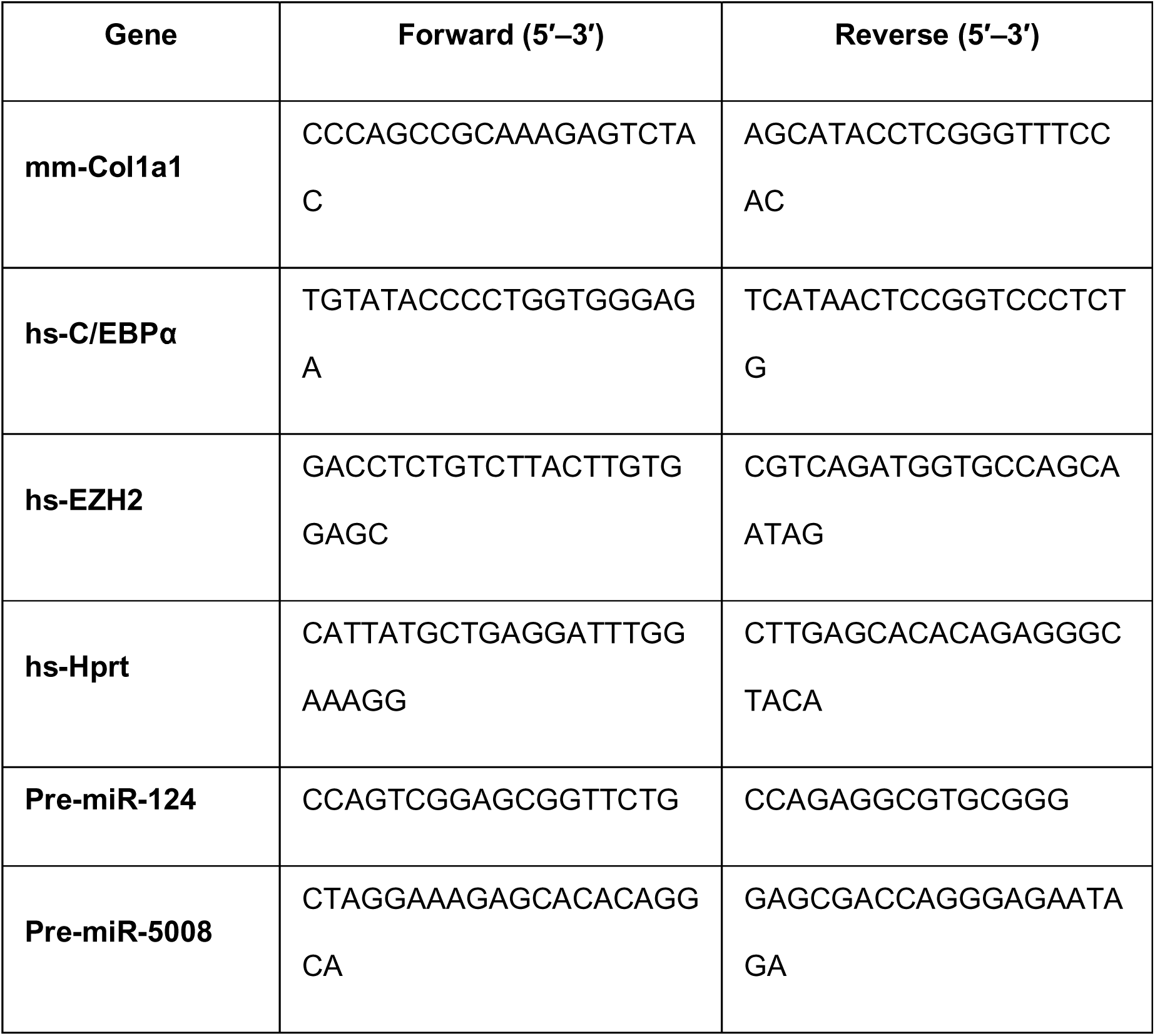
Primer sequences used for qRT-PCR amplification.

### Quantification of mature microRNA levels

miRNA levels were quantified by RT-qPCR using the miRCURY LNA™ RT kit (Qiagen, #339340) for polyadenilation and reverse transcription of the miRNAs. An aliquot of the retrotranscription reaction was quantified with the miRCURY LNA SYBR Green PCR kit (Qiagen #339345) in combination with a specific miRCURY LNA miRNA PCR assay for each miRNA of interest (Qiagen, #339306) or for the reference gene 5S (Qiagen #YP00203906-339306) on a CFX96 Real-Time System (Bio-Rad) as per manufacturer instruction.

### In situ hybridization (ISH) for microRNAs

microRNA in situ hybridization was performed on formalin-fixed paraffin-embedded (FFPE) lung sections (5 µm) using the miRCURY LNA™ microRNA ISH Optimization Kit (Qiagen, # 339451), following the manufacturer’s protocol. Sections were deparaffinized, rehydrated, and digested with proteinase K prior to hybridization with DIG-labeled LNA™ probes (Qiagen, 339115) specific for the target microRNAs. U6 snRNA served as an internal housekeeping control. Hybridization was carried out overnight at 62 °C, followed by stringent washes and incubation with anti-DIG alkaline phosphatase–conjugated antibody (Qiagen, #339451). Signals were detected using NBT/BCIP substrate and counterstained with Nuclear Fast Red.

### Masson’s Trichrome Staining

Masson’s trichrome staining was performed using reagents from Bio-Optica (Milan, Italy). Lung tissues were fixed overnight in 4% PFA at 4 °C, dehydrated in 70% ethanol, embedded in paraffin, and cut into 5 µm sections. Sections were deparaffinized at 60 °C for 2 h, incubated in xylene (30 min) and X-Free solvent (#21-A1305, 5 min), rehydrated through graded ethanols (100%, 90%, 75%, 50%), and washed. Sequential staining was performed with Mayer’s hematoxylin (#21-A06002; 20 min) and Bio-Optica reagents C–F (4–10 min each). Finally, slides were dehydrated, cleared, and mounted in Eukitt medium (Fluka, #03989).

### Hydroxyproline assay

Hydroxyproline content was determined using the Hydroxyproline Assay Kit (Sigma-Aldrich) following the manufacturer’s protocol. Briefly, 10 mg wet lung tissue was homogenized in a MagNA Lyser Instrument (Roche #03 358 968 001) and hydrolyzed in 100 µL sterile water + 100 µL 12 M HCl at 120 °C for 3 h. Samples were centrifuged (10,000 × g, 3 min), and 10 µL supernatant was evaporated at 60 °C. Standards (0.2–1 µg per well) were prepared from a 0.1 mg mL⁻¹ stock. Absorbance was read at 560 nm (microplate reader), and hydroxyproline concentration was calculated from the standard curve.

### Flow cytometry

Lung tissues were processed as described above. Cell suspensions were blocked with FACS BLOCK (BD Biosciences) for 10 min on ice, stained with BD Horizon™ Fixable Viability Stain 510 (1:1000), and labeled for 45 min on ice with the following antibodies (1:200 all): CD45 APC/Cy7, CD31 BV421, EpCAM BV650, MHC II Alexa700, SPC Alexa647 (Abcam), and GFP Alexa488 (Abcam). After washing, cells were analyzed on a BD FACS Aria II cytometer. Doublets were excluded (FSC H vs FSC A, FSC W vs FSC A). Live cells were gated as viability⁺ and alveolar epithelial cells as CD45⁻/CD31⁻/EpCAM⁺/SPC⁺. Unstained, single color, and isotype controls were included. Data were acquired and analyzed using FlowJo (BD Biosciences).

### Generation of AAV6.2FF-miR-124 and AAV6.2FF-miR-5008 vectors

Genomic DNA from HUH7 cells was used as template for PCR amplification of the *pre*-*miR*-*5008* locus using GoTaq Flexi Polymerase (Promega, # M8291). The *pre*-*miR*-*124* sequence was subcloned from an Origene plasmid (#SC400060). Amplicons were re-amplified with chimeric primers containing restriction sites (*Eco*RI/*Not*I for miR-5008 and *Eco*RI/*Eco*RV-blunted for miR-124). All primer sequences are reported in table 4. The pZAC vector was linearized with the corresponding enzymes and ligated to the purified PCR product via Gibson assembly. DH5α competent cells were transformed, and plasmid DNA was purified using the Wizard Plus SV Minipreps Kit (Promega, #A1330). Recombinant plasmids were used for subsequent AAV6.2FF-miR-5008 and AAV6.2FF-miR-124 vector production.

**Table 4.**
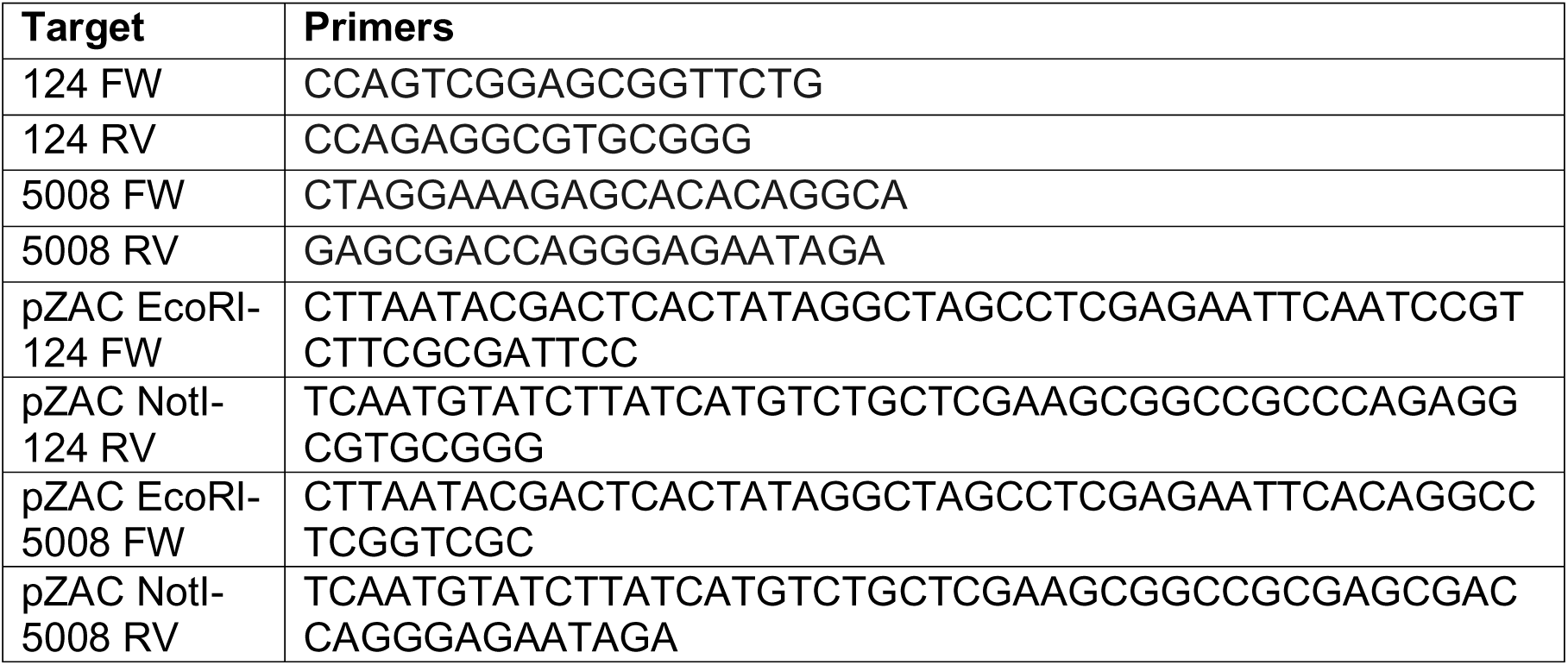
Primer sequences used for genenomic amplification ad Gibson cloning of selected microRNAs.

### AAV vector Preparation

AAV6.2FF-miR-124 and AAV6.2FF-miR-5008 vectors were produced by the AAV Vector Unit at the International Centre for Genetic Engineering and Biotechnology (ICGEB), Trieste (https://www.icgeb.org/avu-core-facility/). Briefly, recombinant AAV particles were generated in HEK293T cells cultured in roller bottles using a cross-packaging strategy, in which the vector genome was packaged into the engineered AAV6.2FF capsid (AAV serotype 6 carrying three additional point mutations), as previously described by Van Lieshout et al. ^32^ Viral vectors were purified by polyethylene glycol (PEG) precipitation followed by cesium chloride (CsCl₂) density gradient centrifugation. The physical titer of the recombinant AAV preparations was determined by quantifying vector genomes encapsidated within viral particles using quantitative real-time PCR against a standard curve.

### Experiments involving the use of animals

All animal experiments were conducted in compliance with the European guidelines and international laws and policies, with the approval of ICGEB Animal Welfare Board, Ethical Committee, and Italian Ministry of Health (Authorizations n° 11/2024-PR (prot. D441B.59) and n. 811/2025-PR (Risp. a prot. D441B.72). Adult male *C57BL/6*, *Sftpc*-*CreERT2*, and *ROSAmTmG*mice (8 weeks old) were used. Mice were randomly assigned into five groups (n = 12 per group): Saline, Bleomycin + AAV6.2FF-GFP, Bleomycin + Nintedanib, Bleomycin + AAV6.2FF-miR-124, Bleomycin + AAV6.2FF-miR-5008. All animals were kept under standard housing conditions (12 h light/dark cycle, controlled temperature). In the prophylactic protocol 30µl of AAV6.2FF constructs (3 × 10¹¹ viral genomes) + 18 µL bleomycin 1U/kg + 12 µL saline were administered simultaneously at day 0. In the therapeutic protocol, 18 µL bleomycin 1U/kg and 42µl saline were administered at D0 and after 10 days from bleomycin injection 30µl of AAV6.2FF constructs (3 × 10¹¹ viral genomes) with 30µl of saline (Vf=60µl) were administered. Nintedanib (60 mg kg⁻¹;) was administered by oral gavage twice daily (b.i.d.) from day 10 to 20 post-bleomycin. At day 21, mice were euthanized, and lungs, livers, and kidneys were collected for downstream analyses.

### Bulk mRNA seq in A549

To identify potential targets of miR-124-3p, A549 cells were transfected with hsa-miR-124-3p. Cells transfected with cel-miR-231-3p (MC4) served as the negative control. Transfections were performed using Lipofectamine RNAiMAX Transfection Reagent (Thermo Fisher Scientific, #13778030) according to the manufacturer’s instructions, with a final microRNA concentration of 25 nM. Briefly, for each well of a 6-well plate, 125 μL of 500 nM microRNA solution was mixed with 125 μL of RNAiMAX transfection mix (117.5 μL Opti-MEM and 7.5 μL RNAiMAX) and incubated for 30 min at 37°C. The resulting complexes were then added dropwise to A549 cells that had been seeded the previous day at a density of 3 × 10^5 cells per well. Three independent biological transfection experiments were performed.Forty-eight hours after transfection, total RNA was isolated from the cells. The culture medium was removed, and each well was lysed with 1 mL of QIAzol Lysis Reagent. RNA was purified using the miRNeasy Mini Kit (Qiagen, #217004) according to the manufacturer’s protocol. Purified RNA was quantified using a NanoDrop spectrophotometer, RNA integrity was assessed by RIN analysis to confirm suitability for sequencing, and high-quality RNA samples were subsequently subjected to RNA sequencing (RNA-seq).

### Library Preparation and RNA Sequencing

RNA concentration and purity were assessed using a NanoDrop ND-1000 spectrophotometer (Thermo Fisher Scientific). RNA integrity and degradation were evaluated using the Agilent 4200 TapeStation system (Agilent Technologies) with the RNA ScreenTape Analysis kit. Only samples with an RNA Integrity Number equivalent (RINe) > 7.0 were processed for downstream sequencing library preparation. Sequencing libraries were prepared using the Illumina TruSeq Stranded mRNA Library Prep Kit (Illumina, San Diego, CA, USA) following the manufacturer’s protocol. Briefly, poly(A)-containing mRNA molecules were purified from 1 µg of total RNA using poly-T oligo-attached magnetic beads. mRNA was fragmented under elevated temperature and prime-transcribed into first-strand cDNA using reverse transcriptase and random hexamers. Second-strand cDNA synthesis was subsequently performed using dUTP to preserve strand specificity. The synthesized cDNA fragments underwent end-repair, 3’ adenylation, and ligation of Illumina sequencing adapters containing unique dual indexes (UDI). The adapter-ligated cDNA library was PCR amplified and purified using AMPure XP beads (Beckman Coulter). Library quality, size distribution, and concentration were validated using the Agilent TapeStation system (D1000 ScreenTape). Prior to pooling, libraries were precisely quantified by quantitative PCR (qPCR) using a KAPA Library Quantification Kit (Roche) to ensure accurate equimolar multiplexing. Quantified libraries were pooled and subjected to paired-end (PE) sequencing 2x150bp on an Illumina NovaSeq 6000 platform. Raw sequencing data were demultiplexed and converted to FASTQ format using the latest version of bcl2fastq2 Conversion Software (Illumina). Raw sequencing data (FASTQ files) were inspected for read quality, adapter contamination, and base quality distribution using FastQC (v0.11.9). Low-quality bases (Phred score (Q < 20) and remaining adapter sequences were trimmed using Cutadapt (v3.4). High-quality trimmed reads were aligned to the reference genome GRCh38 using STAR software. Gene-level read counts were subsequently quantified using featureCounts (Rsubread package) based on the official GENCODE annotation GTF file. All raw sequencing data generated during this study have been deposited in the NCBI Sequence Read Archive (SRA) under the submission ID Sub16377109.

### Pathway analysis

Canonical pathway analysis (IPA) onDEG genes from bulk RNAseq data. Canonical pathway enrichment was performed in Ingenuity Pathway Analysis (IPA, QIAGEN Digital Insights) using as input a filtered list of A549 DEGs from bulk RNAseq data (adjusted *P* < 0.05, log2FC ≥ 0.5 (upregulated) and log2FC ≤ −0.5 (downregulated)), and all genes in the IPA knowledge base as background.

### Differential expression analysis of public scRNA seq dataset

Human single-cell RNA-seq public dataset were analyzed in BioTuring [Le, T., et al. (2020). doi:10.1101/2020.12.11.414136v1]. For each dataset (Habermann/GSE135893 ^7^, Reyfman/GSE122960 ^54^, Yao/GSE146981 ^55^, and Morse/GSE128033 ^56^), AT2 cells were identified according to the annotations provided by the original authors and imported into BBrowserX (BioTuring Browser). Within each dataset, AT2 cells were aggregated to pseudobulk profiles at the subject level (sample-level metadata: Subject ID), and differential expression was performed between AT2 cells from IPF donors (Group 1) and AT2 cells from healthy donors (Group 2) using the pseudobulk DESeq2 method implemented in BBrowserX. For each dataset, genes with false discovery rate (FDR) < 0.05 were selected.

### Bioinformatic analysis to Identify conserved targets of miR-124-3p

To identify hsa-miR-124-3p targets with conserved seed matches, we retrieved predicted target genes from TargetScan Human and TargetScan Mouse (conserved site classes) and restricted the list to genes present in both species, obtaining a list of 1,289 conserved targets. This list was intersected with the set of A549 DEGs (adjusted *P* < 0.05) from the bulk RNA-seq analysis, resulting in 690 conserved targets that were significantly modulated upon hsa-miR-124-3p overexpression in A549 cells.

### Heatmap and Hierarchical clustering analysis

All comparisons of gene expression profiles across datasets were performed in the R environment using the *ComplexHeatmap* package. Gene expression values were visualized as heatmaps, with genes arranged on rows and datasets on columns. Prior to visualization, genes detected in fewer than two public datasets were excluded from the analysis. Hierarchical clustering of genes was performed using Euclidean distance and complete linkage, while dataset ordering was fixed to allow direct comparison between the A549 cell line and primary AT2-derived datasets. Color scales were defined using the *circlize* package to ensure a symmetric representation of expression levels. All graphical outputs were generated in R.

### Statistical Analysis

Statistical analyses were performed using GraphPad Prism (GraphPad Software, San Diego, CA, USA). Data are presented as the mean ± standard deviation (SD). Individual data points are shown in all graphs and represent either biological replicates for in vitro experiments or individual animals for in vivo experiments. Comparisons involving multiple groups were analyzed using one-way or two-way analysis of variance (ANOVA), followed by the appropriate post hoc multiple-comparisons test. Dunnett’s multiple comparisons test was used when comparing multiple experimental groups with a single control, whereas Šídák’s multiple comparisons test was used for selected pairwise comparisons. Where only two groups were compared, statistical significance was determined using an unpaired two-tailed Welch’s *t*-test. Significant values are indicated by the asterisks above the graphs (* P < 0.05,** P < 0,01, *** P < 0,001, **** P < 0,0001).

## Author contribution

L.B. conceived the study. L.B., R.K., and V.M.C. performed the functional screening. V.M.C. and G.Z. performed primary cell isolations, in vitro cellular experiments, and in vivo experiments. L.Z. prepared the AAV vectors. D.L. performed RNAseq on A549 cells, A.M.D.I. performed gene expression analysis and bioinformatic analyses. L.B., G.Z., and V.M.C. analyzed the data. L.B. and G.Z. wrote the original draft of the manuscript. M.V.C., M.C., P.C., and F.S. reviewed and edited the manuscript. All authors read and approved the final manuscript.

## Acknowledgments.

We thank Prof. Serena Zacchigna for generously providing the ROSAmT/mG mice. We are grateful to Dr. Federica Benvenuti’s group, in particular Dr. Giulia Piperno, and Dr. Lucia Ines Lopez Rodriguez, for their technical assistance with flow cytometry. We also thank Ms. Marina Dapas for her technical assistance with AAV vector preparation. We thank Dr. Eszter Nagy, Dr. Gabor Nagy and Dr. Valeria Szijarto for their support of this project through CEBINA GmbH. Graphical Abstract was created in BioRender. Braga, L. (2026) https://BioRender.com/ymi7xdl.

## Declaration of generative AI and AI-assisted technologies in the manuscript preparation process

During the preparation of this work, the authors used ChatGPT Edu (OpenAI) to assist with English language editing and to improve the clarity and readability of the manuscript. The authors carefully reviewed and edited all AI-generated suggestions as needed and take full responsibility for the content of the published article

## Supplementary Figures

**Supplementary Figure 1.**
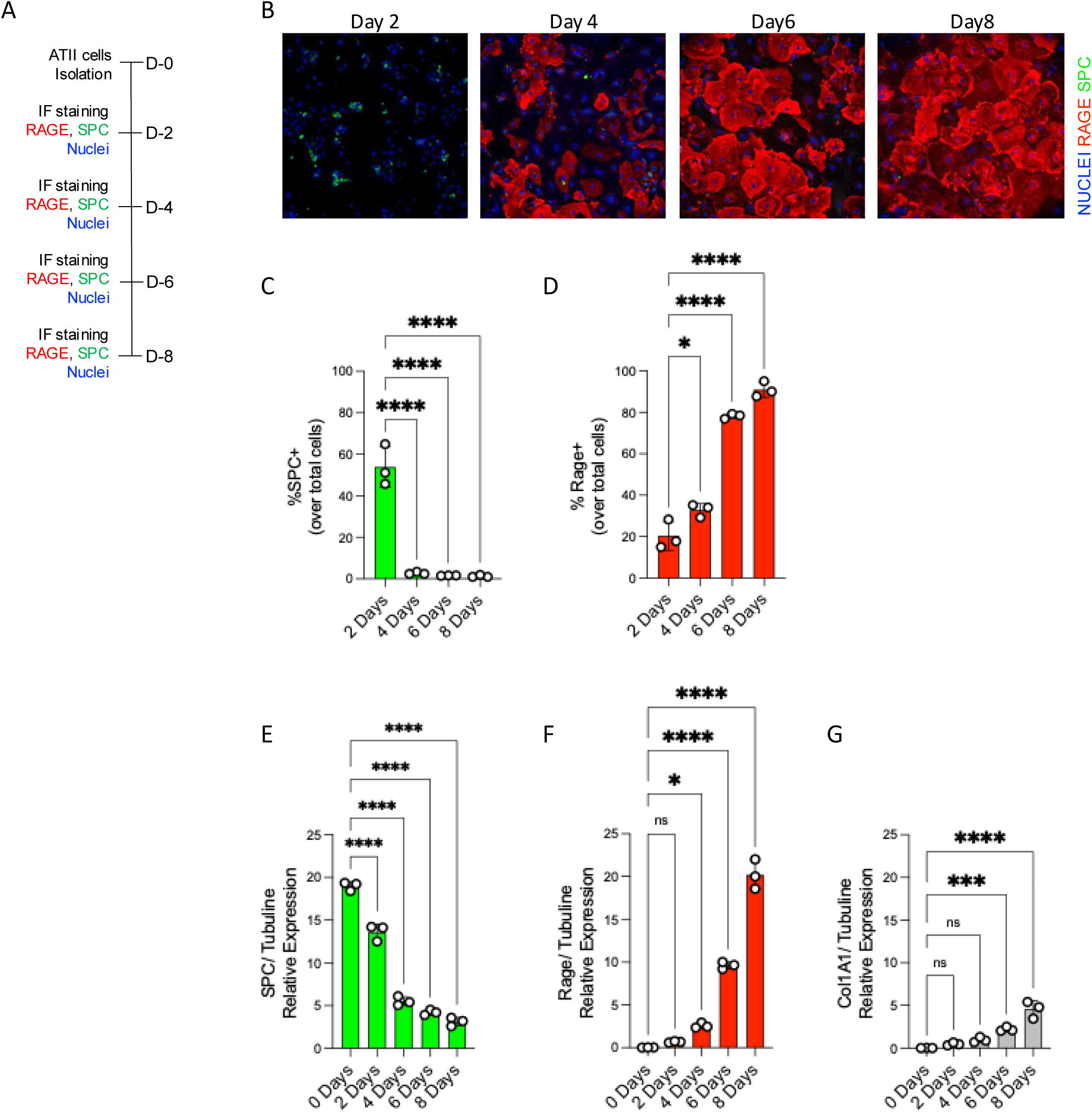
Primary ATII cells isolated from young mice spontaneously transdifferentiate into ATI cells in vitro. (A) Schematic representation of the experimental design. Primary ATII cells were isolated from 8-week-old mice, cultured in vitro, and fixed at 2, 4, 6, and 8 days after plating. (B) Representative immunofluorescence images of primary mouse ATII cells at the indicated time points. Pro-surfactant protein C (pro-SPC, green) was used as a marker of ATII cells, RAGE (red) as a marker of ATI cells, and nuclei were counterstained with Hoechst (blue). (C) Quantitative image analysis of the percentage of pro-SPC–positive cells over total cells. (D) Quantitative image analysis of the percentage of RAGE-positive cells over total cells. E) Real-time PCR quantification of *Sftpc* expression normalized to tubulin. (F) Real-time PCR quantification of *Ager* expression normalized to tubulin. (G) Real-time PCR quantification of *Col1a1* expression normalized to tubulin. Statistical significance was determined by one-way ANOVA followed by Dunnett’s multiple comparisons test. *P < 0.05; **P < 0.01; ***P < 0.0001.

**Supplementary Figure 2.**
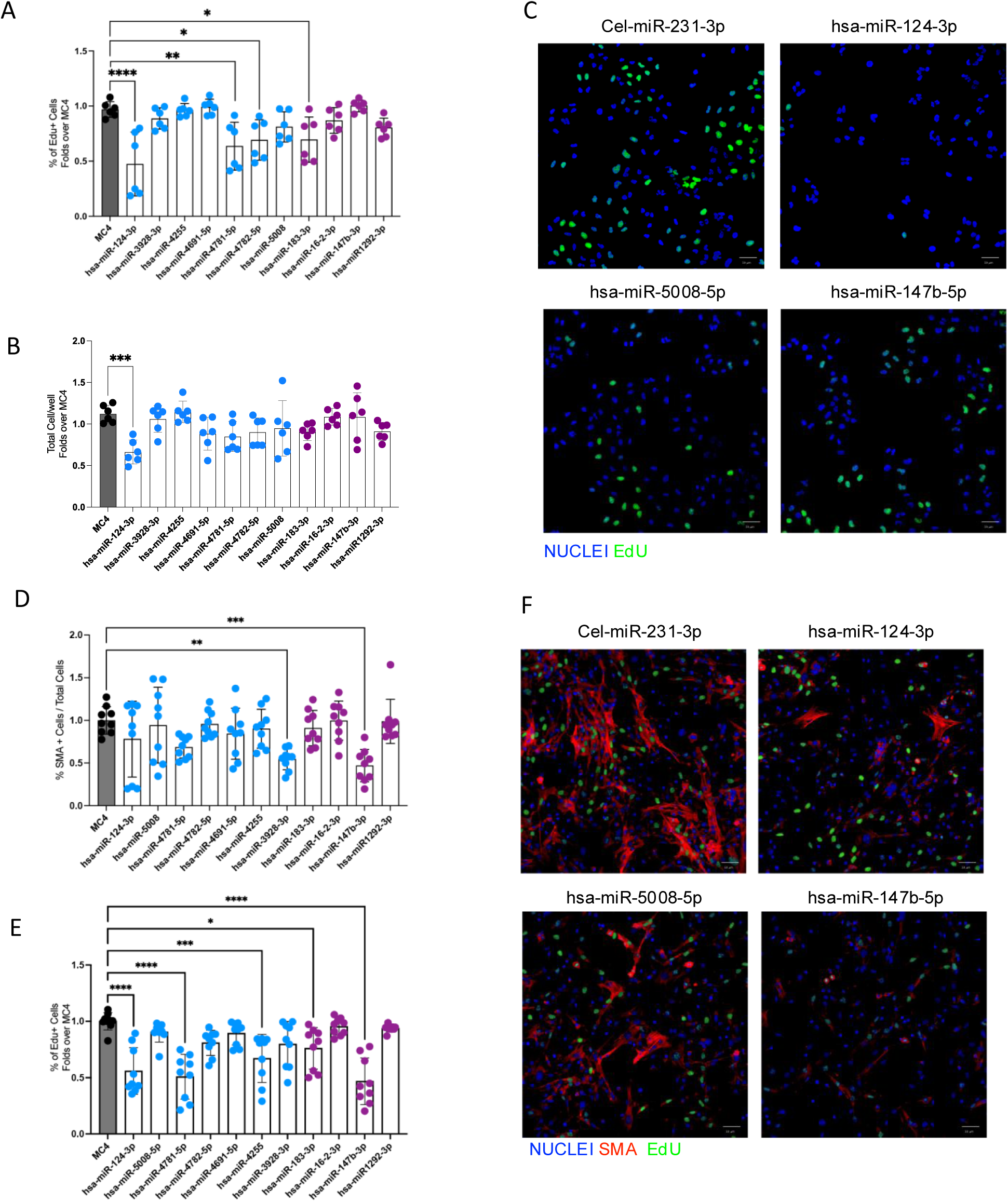
Effects of selected miRNAs on cell proliferation and fibroblast activation. (A) A549 cells were reverse-transfected with selected miRNAs (final concentration, 50 nM) previously identified as promoting (blue dots) or inhibiting (violet dots) ATII-to-ATI transdifferentiation. cel-miR-231-3p (MC4) was used as a negative control. Quantification of EdU-positive cells relative to total cell number is shown as fold change versus MC4. (B) Quantification of total cell number per well in A549 cells treated as in (A). (C) Representative immunofluorescence images of A549 cells treated as in (A). EdU-positive cells are shown in green and nuclei were counterstained with Hoechst (blue). (D) Primary murine lung fibroblasts were reverse-transfected with the same miRNAs and under the same experimental conditions described in (A). Quantitative image analysis of α-SMA–positive cells relative to total cell number is shown as fold change versus MC4. (E) Quantitative image analysis of EdU-positive cells relative to total cell number, expressed as fold change versus MC4, in primary murine lung fibroblasts treated as in (A). (F) Representative immunofluorescence images of primary murine lung fibroblasts treated as in (A). EdU staining (green) identifies proliferating cells, α-SMA staining (red) identifies myofibroblasts, and nuclei were counterstained with Hoechst (blue). Statistical significance was determined by one-way ANOVA followed by Dunnett’s multiple comparisons test. *P < 0.05; **P < 0.01; ***P < 0.0001.

**Supplementary Figure 2.**
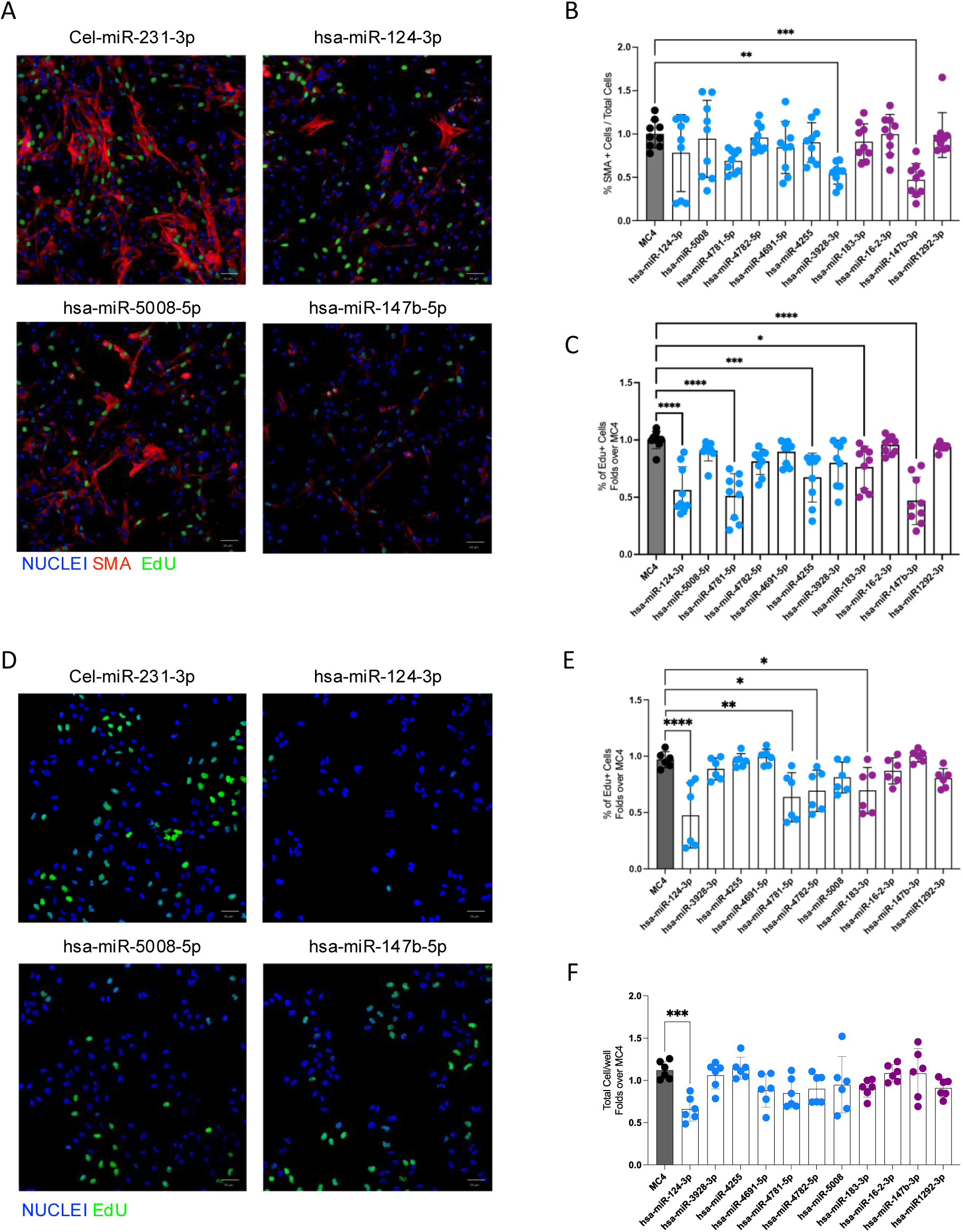
Effects of selected miRNAs on cell proliferation and fibroblast activation. Primary murine lung fibroblasts were reverse-transfected with selected miRNAs (final concentration: 50 nM) previously identified as either promoting (blue dots) or inhibiting (violet dots) ATII-to-ATI transdifferentiation. cel-miR-231-3p (MC4) was used as a negative control. (**A**) Representative immunofluorescence images of primary murine lung fibroblasts stained for α-SMA (red), EdU (green), and nuclei (blue) following treatment with the indicated miRNAs. (**B**) Quantitative image analysis of α-SMA-positive primary murine lung fibroblasts relative to the total cell number, expressed as fold change compared with MC4. (**C**) Quantitative image analysis of EdU-positive primary murine lung fibroblasts relative to the total cell number, expressed as fold change compared with MC4. (**D**) Representative immunofluorescence images of A549 cells reverse-transfected with the indicated miRNAs (50 nM) and stained for EdU (green) and nuclei (blue). cel-miR-231-3p (MC4) was used as a negative control. (**E**) Quantitative image analysis of the total number of A549 cells, expressed as fold change compared with MC4. (**F**) Quantitative image analysis of EdU-positive A549 cells relative to the total cell number, expressed as fold change compared with MC4. Statistical significance was determined by one-way ANOVA followed by Dunnett’s multiple comparisons test. *P < 0.05; **P < 0.01; ***P < 0.0001.

**Supplementary Figure 3.**
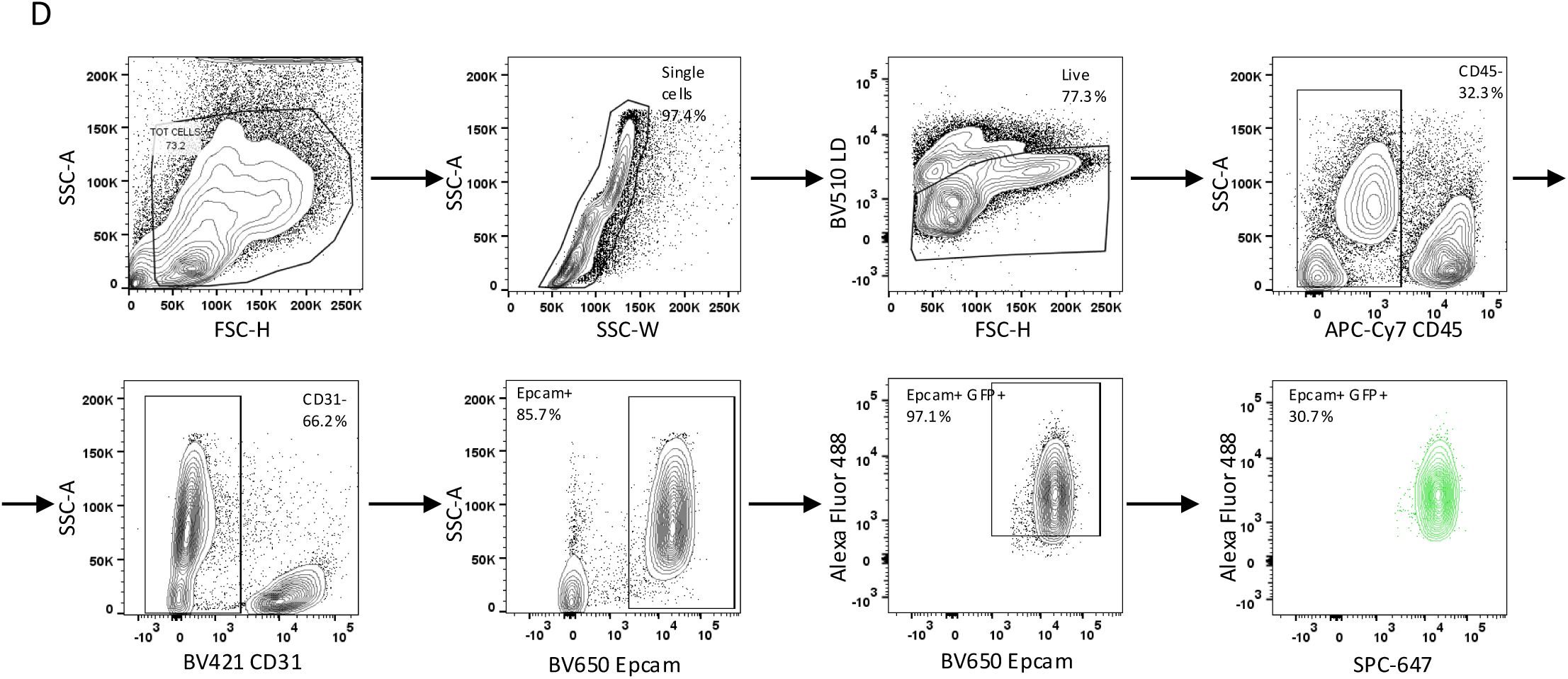
Flow cytometry gating strategy for the quantification of GFP-positive ATII cells 7 days after intratracheal injection of AAV6.2FF-GFP. Total lung tissue was enzymatically digested and filtered through a 200 μm mesh. Cells were stained with Live/Dead dye and antibodies against CD45, CD31, EpCAM, pro-SPC, and GFP. The gating strategy was performed as follows: first, debris and doublets were excluded based on forward and side scatter parameters to select single cells. Live cells were then identified by exclusion of dead cells. Within the live cell population, CD45-positive cells (immune cells) were excluded, followed by exclusion of CD31-positive cells (endothelial cells). Among the live CD45⁻/CD31⁻ population, EpCAM-positive epithelial cells were selected. Within EpCAM-positive cells, GFP-positive cells were identified. Finally, the percentage of pro-SPC–positive cells was quantified within the EpCAM⁺/GFP⁺ population. Using this strategy, approximately 30% of ATII cells were identified as GFP-positive, indicating successful transduction by AAV6.2FF-GFP.

**Supplementary Figure 4.**
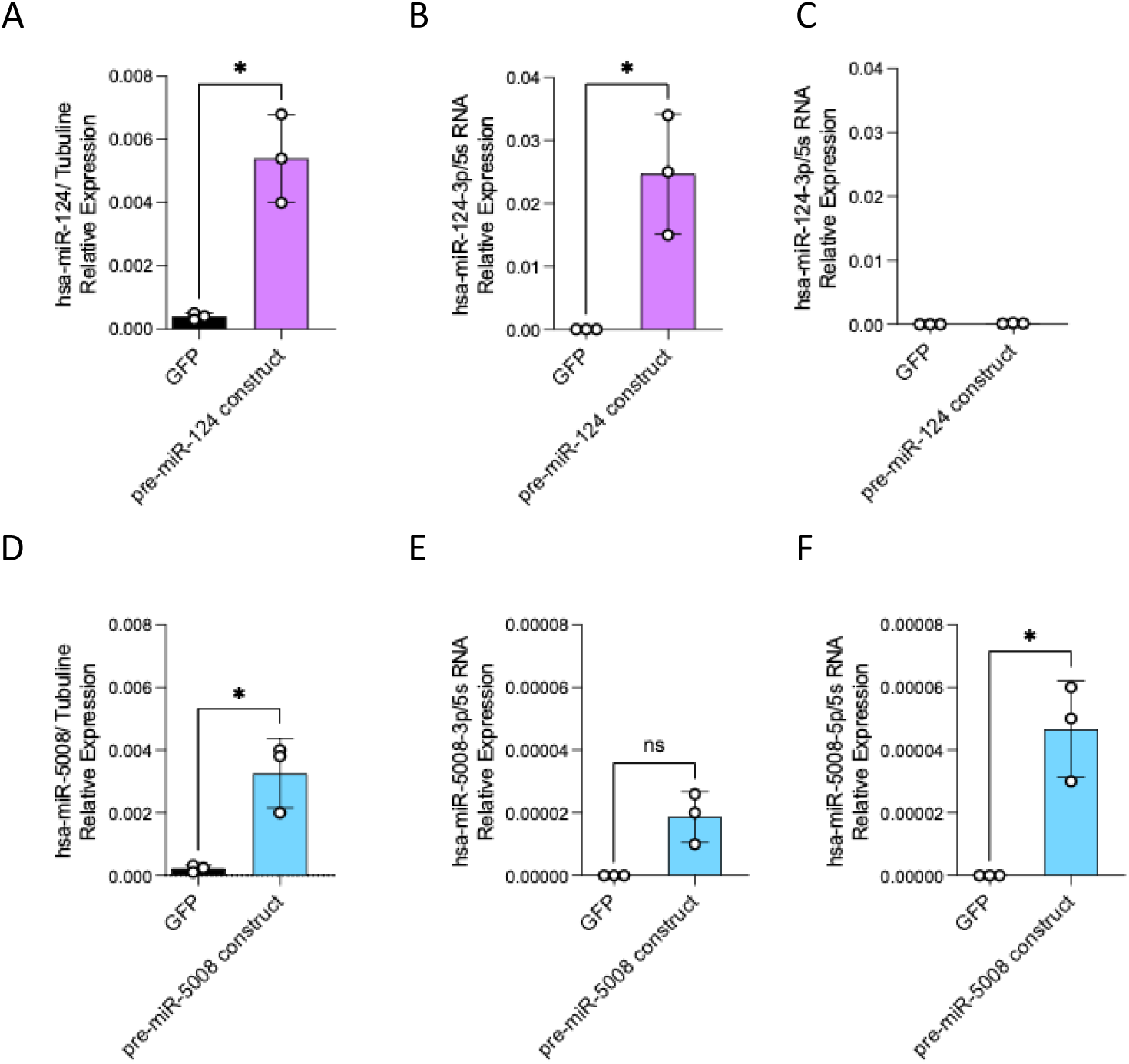
AAV6.2FF-mediated delivery of miR-124 and miR-5008 and assessment of strand specificity. The efficiency of AAV6.2FF-mediated delivery of miR-124 and miR-5008 and the assessment of strand specificity (3p vs 5p) were evaluated. Mice were intratracheally injected with 30 μL of AAV6.2FF-miRNA vectors (1 × 10¹³ vg/mL). Lungs were harvested 7 days post-injection for RNA analysis. (A) Real-time PCR quantification of pre-miR-124 expression normalized to tubulin. LNA-based quantitative PCR analysis of mature miR-124-3p (B) and miR-124-5p (C) expression in total lung tissue 7 days after intratracheal AAV6.2FF administration. The -3p strand was robustly expressed. (D) Real-time PCR quantification of pre-miR-5008 expression normalized to tubulin. LNA-based quantitative PCR analysis of mature miR-5008-3p (E) and miR-5008-5p (F) expression in total lung tissue 7 days after intratracheal AAV6.2FF administration. Both the -3p and -5p strand was robustly expressed. Statistical significance for pairwise comparisons were analyzed using an unpaired two-tailed Welch’s t-test.. *P < 0.05; **P < 0.01; ***P < 0.0001.

**Supplementary Figure 5.**
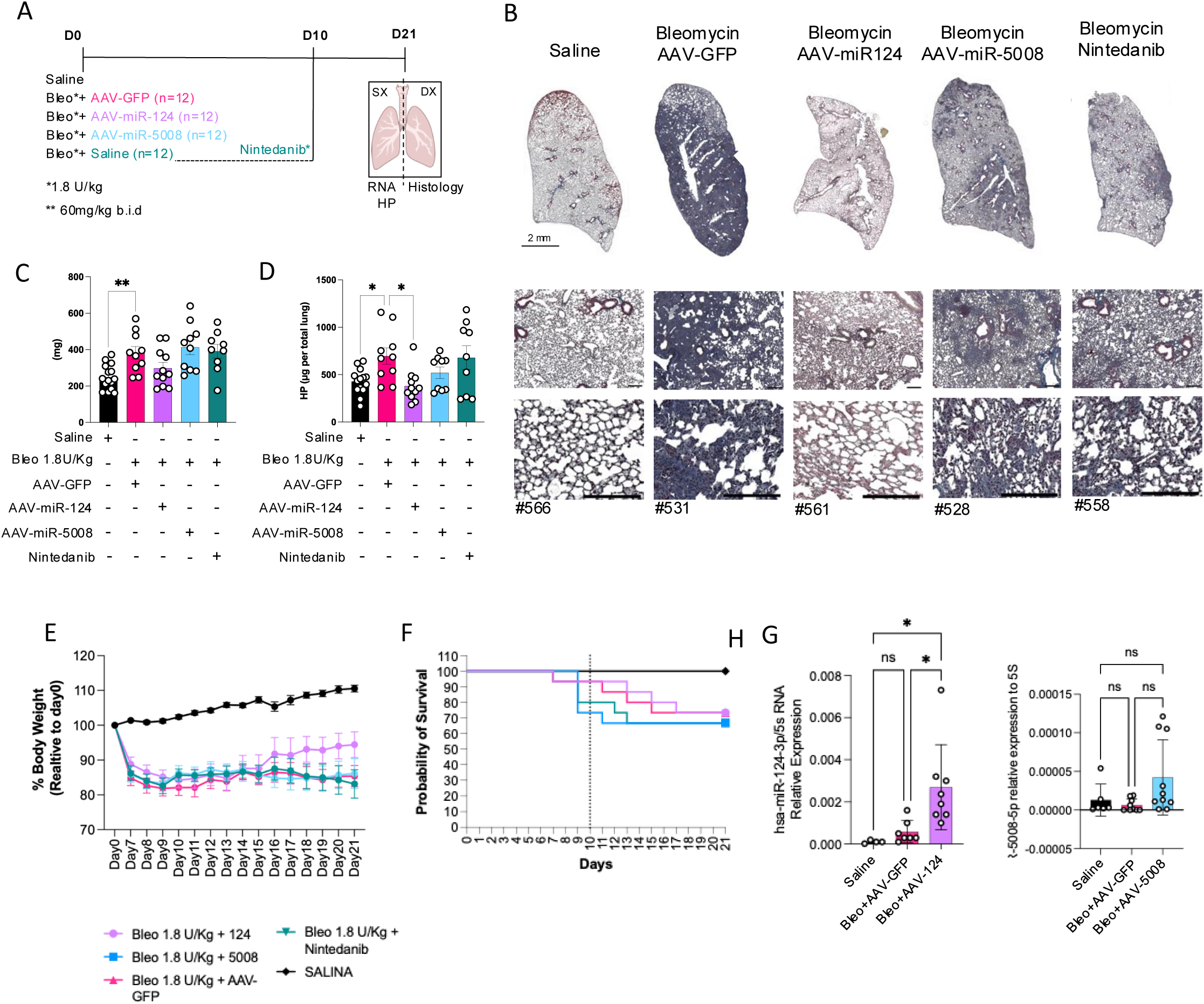
Preventive administration of AAV6.2FF-miR-124 and AAV6.2FF-miR-5008 in bleomycin-induced lung fibrosis. (A) Schematic overview of the experimental design to evaluate the preventive effect of miR-124 and miR-5008 delivered by AAV6.2FF in bleomycin-induced lung fibrosis. Mice received intratracheal bleomycin (1.8U/Kg) alone or in combination with 3x10^11 Vg of AAV6.2FF-GFP (Negative Control) or AAV6.2FF-miR-124 or AAV6.2FF-miR-5008. On day 10, a group bleomycin-treated animals received Nintedanib at 60 mg/kg twice daily (BID) via oral gavage . Mice were sacrificed 20 days after bleomycin administration. Lungs were harvested; the right lobe was processed for RNA extraction and hydroxyproline quantification, whereas the left lobe was used for histological analyses (Masson’s trichrome staining and immunofluorescence). (B) Representative Masson’s trichrome staining of lung sections from mice treated as in A. (C) Lung weight and (D) Quantification of lung hydroxyproline content quantification for treatment group as in A. (E) Body weight changes over time relative to day 0. (F) Kaplan–Meier survival analysis. miRCURY LNA miRNA-based quantitative PCR analysis of mature miR-124-3p (G) and miR-5008-5p (H) expression in total lung tissue 20 days after bleomycin administration. Statistical significance was determined by one-way ANOVA followed by Dunnett’s multiple comparisons test. *P < 0.05; **P < 0.01; ***P < 0.0001.

**Supplementary Figure 6.**
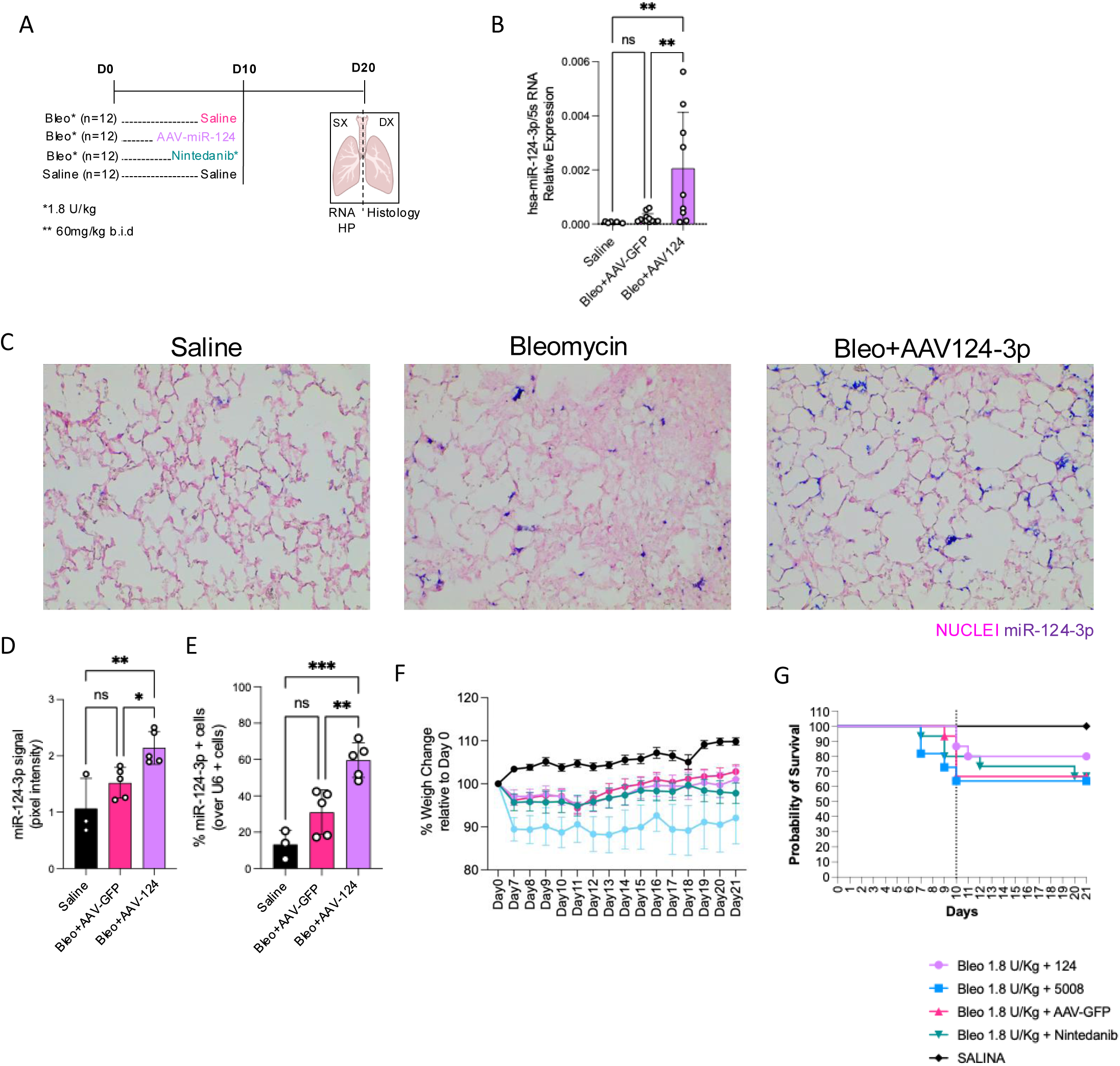
Therapeutic administration of AAV6.2FF-miR-124 in bleomycin-induced lung fibrosis. (A) Schematic overview of the experimental design to evaluate the therapeutic potential of miR-124 delivered by AAV6.2FF in bleomycin-induced lung fibrosis. Mice received intratracheal bleomycin. Ten days later, at the onset of the fibrotic phase, animals were treated with AAV6.2FF-GFP (control), AAV6.2FF-miR-124, or nintedanib. Mice were sacrificed 21 days after bleomycin administration. Lungs were harvested; the right lobe was processed for RNA extraction and hydroxyproline quantification, whereas the left lobe was used for histological analyses (in situ hybridization against miR-124-3p). (B) miRCURY LNA miRNA-based quantitative PCR analysis of mature miR-124-3p expression in total lung tissue 21 days after bleomycin administration for treatmentgroup as in A . (C) Representative in situ hybridization (ISH) staining of lung sections for treatment group as in A. miR-124-3p was detected using a digoxigenin-labeled probe and visualized by alkaline phosphatase (AP)-based chromogenic development (purple signal), while nuclei were counterstained with Sirius Red (pink). Quantification of ISH analysis for treatment group as in A expressed as the (D) Percentage of miR-124-3p–positive cells over total U6-positive cells (miRNA internal control) or (E) miR-124-3p signal intensity over the total lung section area, expressed as pixel intensity. (F) Body weight changes over time relative to day 0. Only mice treated with AAV6.2FF-miR-124 showed recovery of body weight comparable to PBS-treated control mice.(G) Kaplan–Meier survival analysis. Overall survival was approximately 60% across all treatment groups, with no significant differences observed. Statistical significance was determined by one-way ANOVA followed by Dunnett’s multiple comparisons test. *P < 0.05; **P < 0.01; ***P < 0.0001.

**Supplementary Figure 7.**
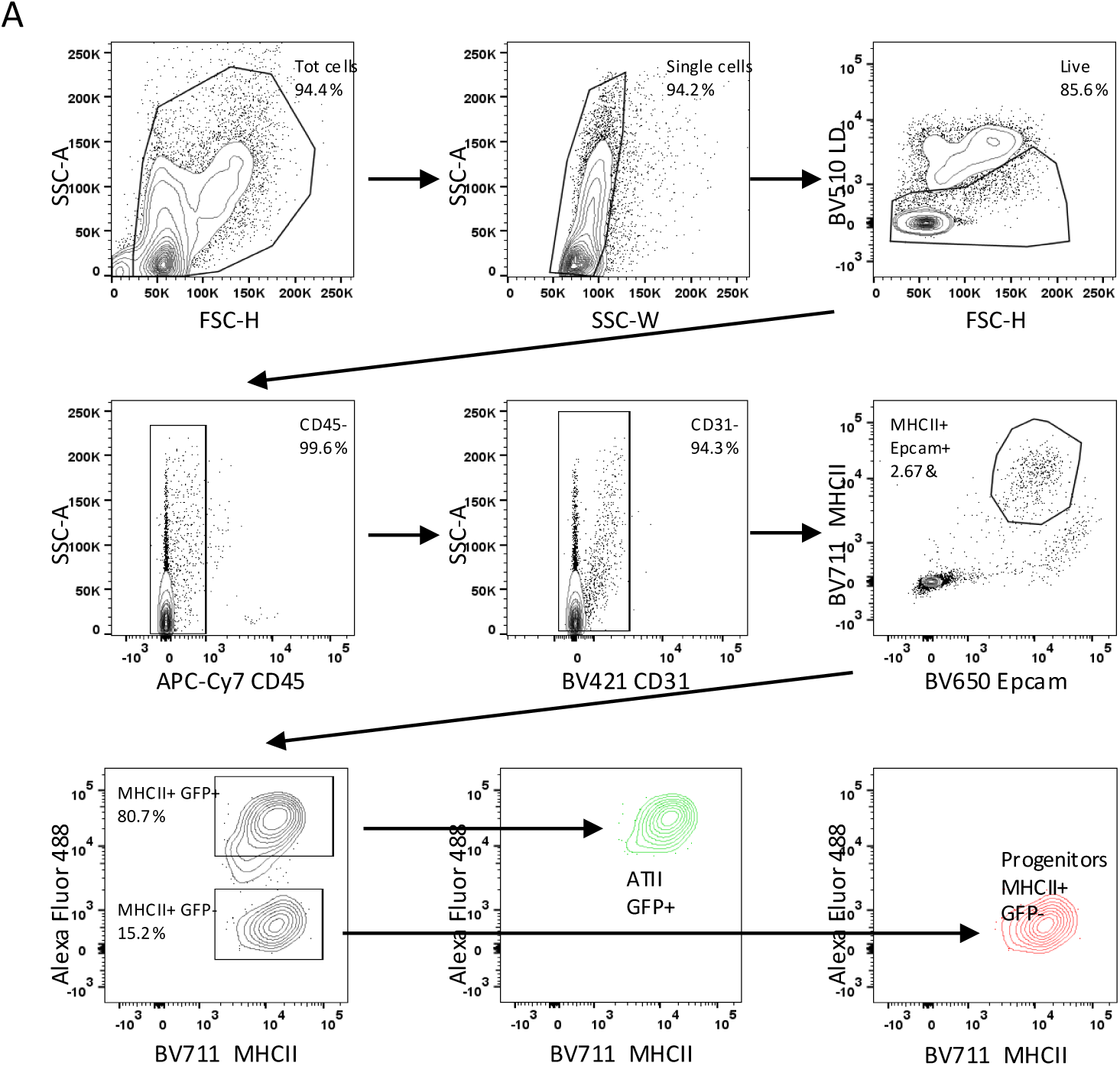
Flow cytometry gating strategy confirming the presence of MHC-II^⁺^/GFP^⁻^ cells in isolated ATII populations. (A) ATII cells were isolated from SFTPC-CreER[T2]/mTmG mice treated with tamoxifen. Cells were stained with a Live/Dead dye and antibodies against CD45, CD31, EpCAM, MHC-II, and GFP. The gating strategy was applied as follows: debris and doublets were excluded based on forward and side scatter parameters to select single cells, and live cells were identified by exclusion of dead cells. Within the live population, CD45⁺ immune cells and CD31⁺ endothelial cells were excluded. Epithelial cells were then selected as EpCAM⁺ cells within the CD45⁻/CD31⁻ population. Among EpCAM⁺ cells, MHC-II⁺ cells were identified, and GFP expression was analyzed to distinguish mature ATII cells (GFP⁺, SPC-expressing) from GFP⁻ cells. The percentages of MHC-II⁺/GFP⁺ cells (mature ATII cells) and MHC-II⁺/GFP⁻ cells (putative ATII progenitor-like cells) were quantified within the EpCAM⁺/MHC-II⁺ gate. Using this approach, the ATII population consisted of approximately 80.7% MHC-II⁺/GFP⁺ mature ATII cells and 15.2% MHC-II⁺/GFP⁻ progenitor-like cells. In the final gating plot, mature ATII cells are shown in green and progenitor-like ATII cells in red.

**Supplementary Figure 8.**
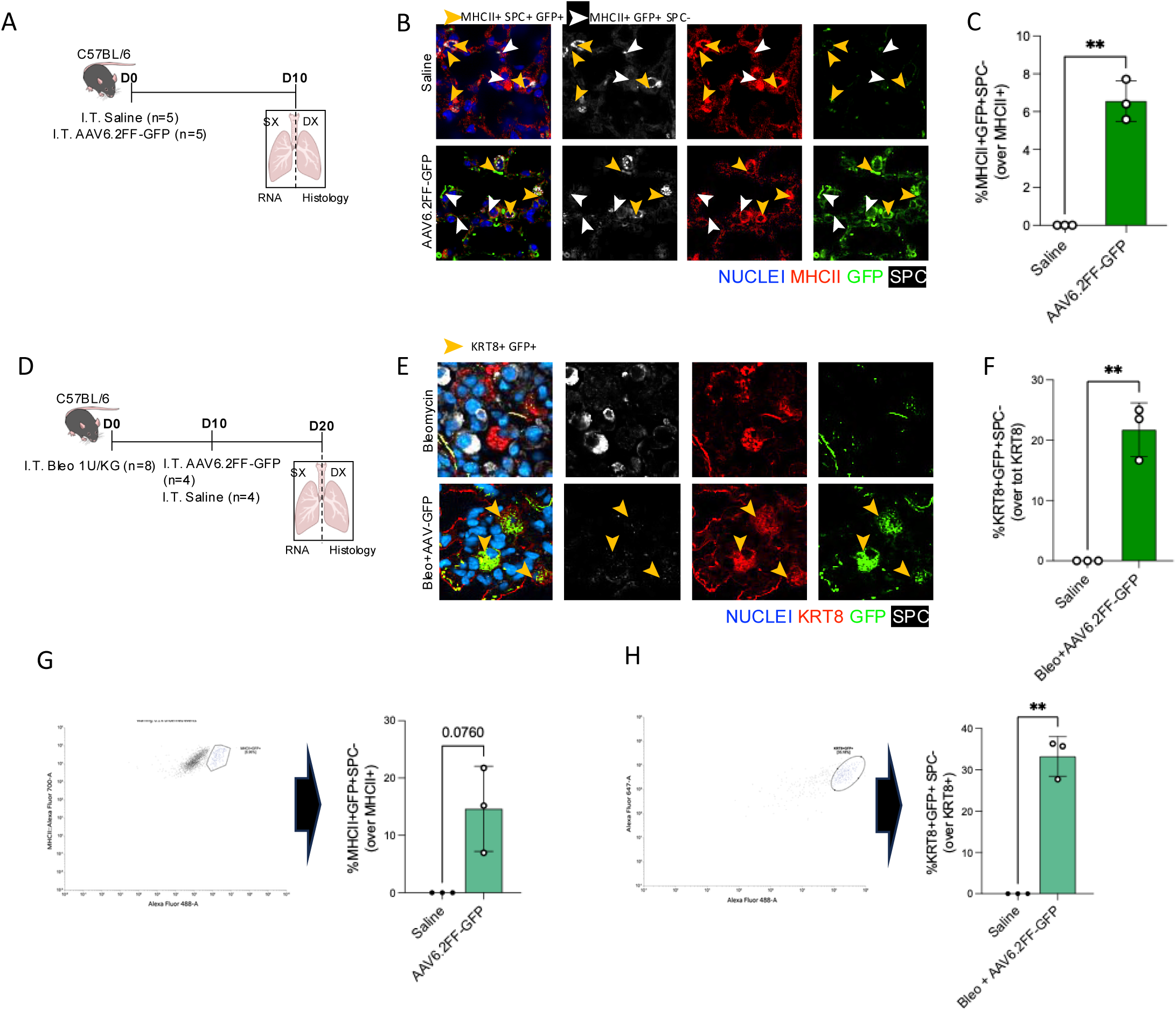
AAV6.2FF transduction efficiency in ATII subpopulations and transitional intermediates. (A) Schematic overview of the experimental workflow to assess the efficacy of AAV6.2FF-mediated transduction in progenitor ATII cells. AAV6.2FF-GFP (30 μL, 1 × 10¹³ vg/mL) was administered by intratracheal injection. Mice were sacrificed 10 days later. Lungs were harvested; the right lobe was processed for RNA extraction, whereas the left lobe was used for histological analysis (immunofluorescence staining). (B) Representative immunofluorescence images of lung sections collected 7 days after intratracheal injection of AAV6.2FF-GFP or saline (control). ATII cells are stained for pro-SPC (white), GFP is shown in green, MHCII (marker of progenitor and mature ATII cells) is shown in red, and nuclei are counterstained with Hoechst (blue). (C) Quantitative image analysis of progenitor-like ATII cells (MHCII⁺/GFP^+^/SPC⁻) transduced by AAV6.2FF-GFP. Approximately 6.67% ±1.1% of progenitor-like ATII cells were GFP-positive. G) Flow Cytometry of progenitors MHCII-club cells transduced by AAV6.2FF-GFP. Singlets were selected followed by exclusion of dead cells and doublets. Lineage-negative cells (CD31⁻CD45⁻) were then gated, and epithelial cells identified as EpCAM⁺. Within this population, MHCII⁺ cells were selected and ATII cells (SPC⁺) were excluded,to identified the MHCII-club progenitor fraction. In this population GFP+ cells were selected (14.64%± 7,4%). (D) Schematic overview of the experimental workflow to assess AAV6.2FF-mediated transduction in aberrant transitional intermediates (KRT8⁺ cells) during bleomycin-induced lung injury. Bleomycin (1 U/kg) was administered intratracheally. Ten days later, AAV6.2FF-GFP (30 μL, 1 × 10¹³ vg/mL) was delivered intratracheally. Mice were sacrificed 20 days after bleomycin injection. Lungs were harvested; the right lobe was processed for RNA extraction, whereas the left lobe was used for immunofluorescence analysis. (E) Representative immunofluorescence images of lung sections collected 20 days after bleomycin injection. ATII cells are stained for pro-SPC (white), GFP is shown in green, KRT8 (marker of aberrant transitional intermediates) is shown in red, and nuclei are counterstained with Hoechst (blue). (F) Quantitative image analysis of aberrant transitional intermediates (KRT8⁺/GFP^+^/SPC⁻) transduced by AAV6.2FF-GFP. Approximately 21.7%±4.5% of KRT8⁺ cells were transduced. (H) Flow Cytometry of KRT8+ cells transduced by AAV6.2FF-GFP. Singlets were selected from total lung dissociates, followed by exclusion of dead cells and doublets. Lineage-negative cells (CD31⁻CD45⁻) were gated, and epithelial cells identified as EpCAM⁺. Within this population, KRT8⁺ cells were selected as aberrant transitional cell fraction. Then GFP positivity within this gate identified 33.23%± 4.8% of KRT8+GFP+ cells. Statistical significance for pairwise comparisons were analyzed using an unpaired two-tailed Welch’s t-test.. *P < 0.05; **P < 0.01; ***P < 0.0001.

**Supplementary Figure 9.**
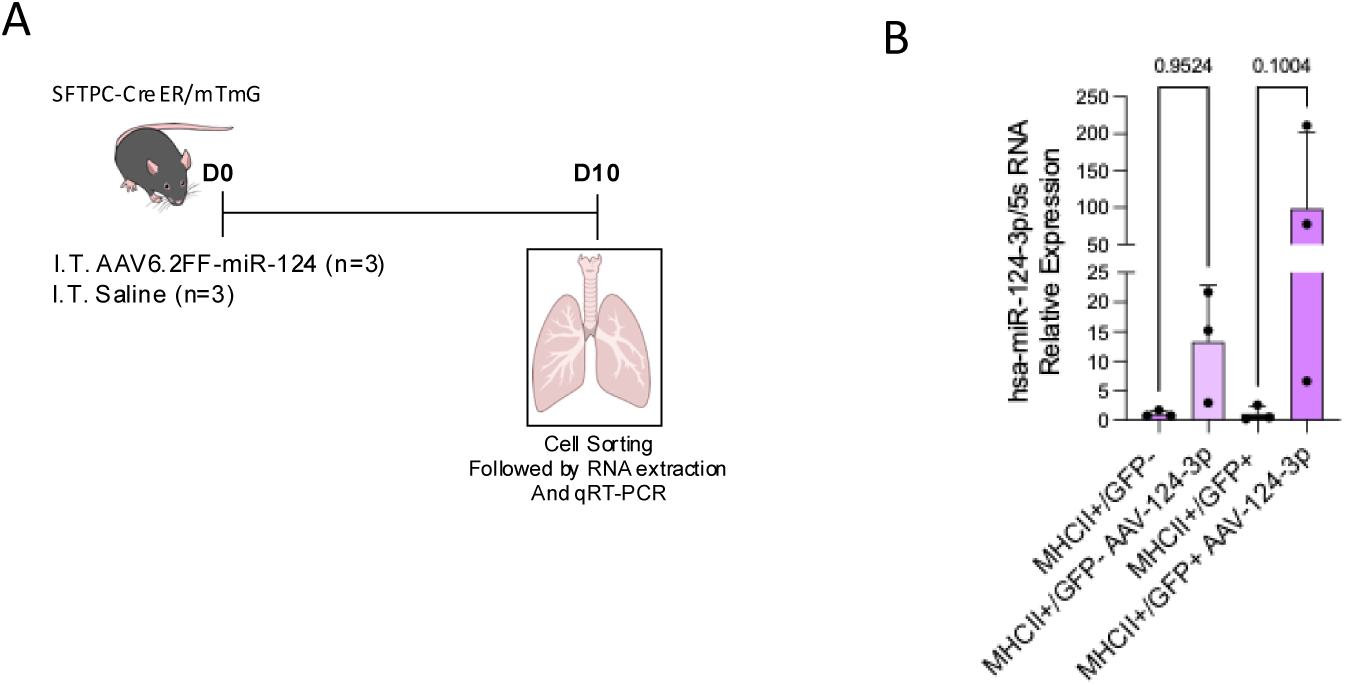
AAV6.2FF driven mIR-124-3p expression in MHC-II^⁺^ Club distal progenitor cells and ATII cells. (A) Schematic overview of the experimental design to evaluate the expression levels of miR-124 in ATII cells (MHC-II+/GFP+) and in progenitors MHC-II^+^ club cells (MHC-II+/GFP-). Mice were injected intratracheally with AAV6.2FF-miR-124(30 μL, 1 × 10¹³ vg/mL), ten days later mice were sacrificed. Cells were sorted based on the gate strategy described previously (A) and RNA was extracted to quantify the expression levels of miR-124-3p. (B) miRCURY LNA miRNA-based quantitative PCR analysis of mature miR-124-3p expression in progenitors (MHC-II+/GFP-) and ATII cells (MHC-II+/GFP+) sorted from saline mice and mice injected with AAV-124-3p. Statistical significance was determined using one-way ANOVA applied to selected pairwise comparisons between specific groups, Šídák’s multiple comparisons test was used. *P < 0.05; **P < 0.01; ***P < 0.0001.

**Supplementary Figure 10.**
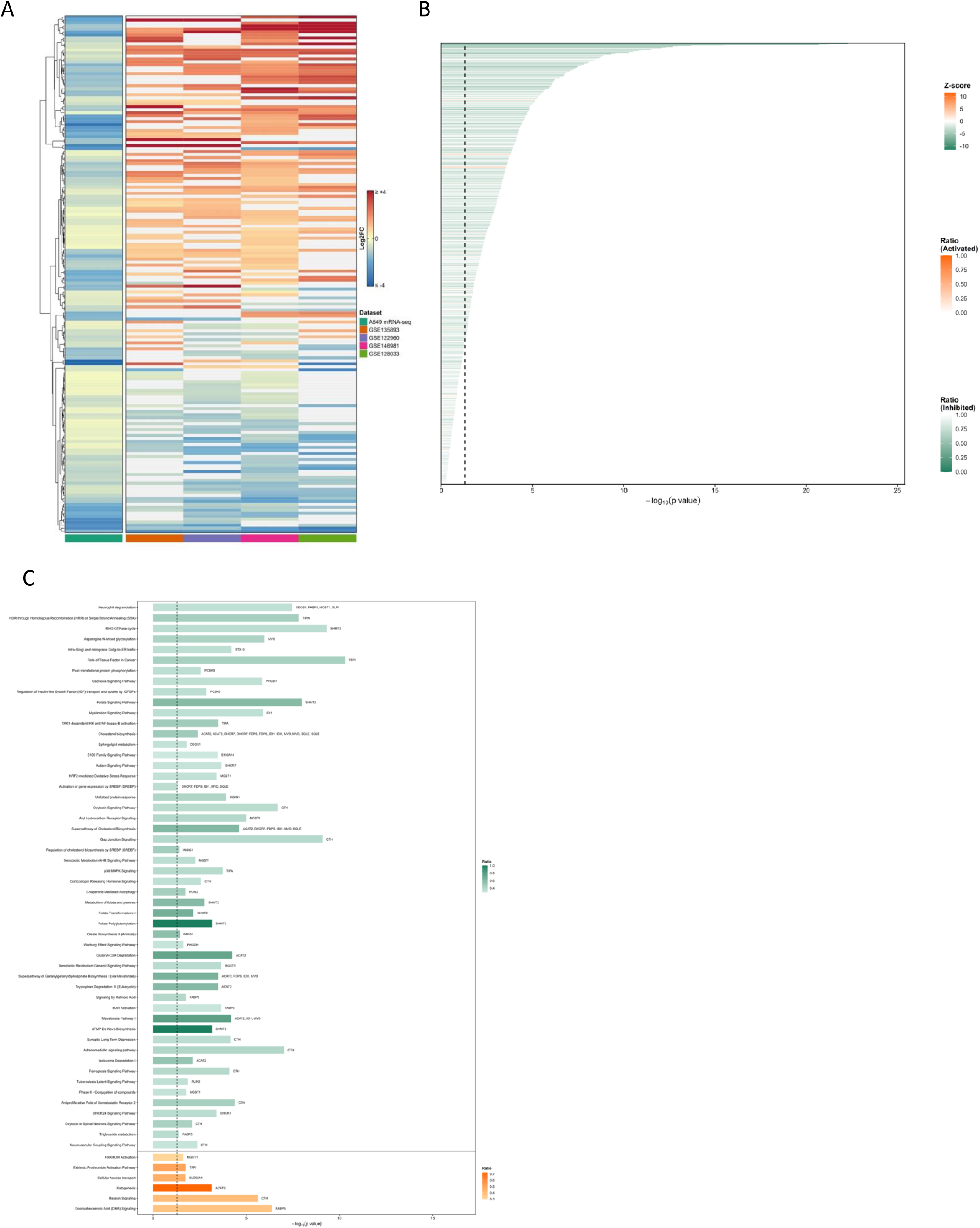
Bioinformatic analysis for the identification of candidate miR-124-3p targets in human disease. (A) Heatmap showing the comparative transcriptomic analysis between genes significantly downregulated in A549 cells following miR-124-3p transfection and differentially expressed genes identified in ATII cells from IPF versus healthy lungs across four independent public single-cell RNA-seq datasets: Habermann/GSE135893, Reyfman/GSE122960, Yao/GSE146981 and Morse/GSE128033. Genes were retained when differential expression data were available in at least two public datasets. Rows represent genes and columns represent datasets; color intensity indicates the relative direction and magnitude of differential expression. (B) Canonical pathway analysis of genes downregulated in A549 cells following miR-124-3p transfection compared with MC4 control, performed using Ingenuity Pathway Analysis (IPA, QIAGEN). Enriched pathways are shown as horizontal bars and ordered according to statistical significance. Predicted pathway activation or inhibition was assigned according to the IPA activation z-score, where available. (C) Horizontal bar plot showing IPA canonical pathway annotation of the shared downregulated gene set identified through the cross-dataset comparison. Enriched pathways are ranked by statistical significance. Predicted pathway activation or inhibition was assigned according to the IPA activation z-score, where available.

## References

1. Barkauskas, C.E., Cronce, M.J., Rackley, C.R., Bowie, E.J., Keene, D.R., Stripp, B.R., Randell, S.H., Noble, P.W., and Hogan, B.L. (2013). Type 2 alveolar cells are stem cells in adult lung. J Clin Invest 123, 3025–3036. 10.1172/JCI68782.

2. Rock, J.R., Barkauskas, C.E., Cronce, M.J., Xue, Y., Harris, J.R., Liang, J., Noble, P.W., and Hogan, B.L. (2011). Multiple stromal populations contribute to pulmonary fibrosis without evidence for epithelial to mesenchymal transition. Proc Natl Acad Sci U S A 108, E1475–1483. 10.1073/pnas.1117988108.

3. Jiang, P., Gil de Rubio, R., Hrycaj, S.M., Gurczynski, S.J., Riemondy, K.A., Moore, B.B., Omary, M.B., Ridge, K.M., and Zemans, R.L. (2020). Ineffectual Type 2-to-Type 1 Alveolar Epithelial Cell Differentiation in Idiopathic Pulmonary Fibrosis: Persistence of the KRT8(hi) Transitional State. Am J Respir Crit Care Med 201, 1443–1447. 10.1164/rccm.201909-1726LE.

4. Raghu, G., Remy-Jardin, M., Myers, J.L., Richeldi, L., Ryerson, C.J., Lederer, D.J., Behr, J., Cottin, V., Danoff, S.K., Morell, F., et al. (2018). Diagnosis of Idiopathic Pulmonary Fibrosis. An Official ATS/ERS/JRS/ALAT Clinical Practice Guideline. Am J Respir Crit Care Med 198, e44–e68. 10.1164/rccm.201807-1255ST.

5. Raghu, G., Remy-Jardin, M., Richeldi, L., Thomson, C.C., Inoue, Y., Johkoh, T., Kreuter, M., Lynch, D.A., Maher, T.M., Martinez, F.J., et al. (2022). Idiopathic Pulmonary Fibrosis (an Update) and Progressive Pulmonary Fibrosis in Adults: An Official ATS/ERS/JRS/ALAT Clinical Practice Guideline. Am J Respir Crit Care Med 205, e18–e47. 10.1164/rccm.202202-0399ST.

6. Selman, M., and Pardo, A. (2020). The leading role of epithelial cells in the pathogenesis of idiopathic pulmonary fibrosis. Cell Signal 66, 109482. 10.1016/j.cellsig.2019.109482.

7. Habermann, A.C., Gutierrez, A.J., Bui, L.T., Yahn, S.L., Winters, N.I., Calvi, C.L., Peter, L., Chung, M.I., Taylor, C.J., Jetter, C., et al. (2020). Single-cell RNA sequencing reveals profibrotic roles of distinct epithelial and mesenchymal lineages in pulmonary fibrosis. Sci Adv 6, eaba1972. 10.1126/sciadv.aba1972.

8. Strunz, M., Simon, L.M., Ansari, M., Kathiriya, J.J., Angelidis, I., Mayr, C.H., Tsidiridis, G., Lange, M., Mattner, L.F., Yee, M., et al. (2020). Alveolar regeneration through a Krt8+ transitional stem cell state that persists in human lung fibrosis. Nat Commun 11, 3559. 10.1038/s41467-020-17358-3.

9. Kim, C.F., Jackson, E.L., Woolfenden, A.E., Lawrence, S., Babar, I., Vogel, S., Crowley, D., Bronson, R.T., and Jacks, T. (2005). Identification of bronchioalveolar stem cells in normal lung and lung cancer. Cell 121, 823–835. 10.1016/j.cell.2005.03.032.

10. Liu, Q., Liu, K., Cui, G., Huang, X., Yao, S., Guo, W., Qin, Z., Li, Y., Yang, R., Pu, W., et al. (2019). Lung regeneration by multipotent stem cells residing at the bronchioalveolar-duct junction. Nat Genet 51, 728–738. 10.1038/s41588-019-0346-6.

11. Liu, K., Meng, X., Liu, Z., Tang, M., Lv, Z., Huang, X., Jin, H., Han, X., Liu, X., Pu, W., et al. (2024). Tracing the origin of alveolar stem cells in lung repair and regeneration. Cell 187, 2428–2445 e2420. 10.1016/j.cell.2024.03.010.

12. Kathiriya, J.J., Brumwell, A.N., Jackson, J.R., Tang, X., and Chapman, H.A. (2020). Distinct Airway Epithelial Stem Cells Hide among Club Cells but Mobilize to Promote Alveolar Regeneration. Cell Stem Cell 26, 346–358 e344. 10.1016/j.stem.2019.12.014.

13. Wang, F., Ting, C., Riemondy, K.A., Douglas, M., Foster, K., Patel, N., Kaku, N., Linsalata, A., Nemzek, J., Varisco, B.M., et al. (2023). Regulation of epithelial transitional states in murine and human pulmonary fibrosis. J Clin Invest 133. 10.1172/JCI165612.

14. Confalonieri, P., Volpe, M.C., Jacob, J., Maiocchi, S., Salton, F., Ruaro, B., Confalonieri, M., and Braga, L. (2022). Regeneration or Repair? The Role of Alveolar Epithelial Cells in the Pathogenesis of Idiopathic Pulmonary Fibrosis (IPF). Cells 11. 10.3390/cells11132095.

15. Hewitt, R.J., Pearmain, L., Lyka, E., and Dickens, J. (2025). Epithelial damage and ageing: the perfect storm. Thorax 80, 668–675. 10.1136/thorax-2024-222060.

16. Bartel, D.P. (2004). MicroRNAs: genomics, biogenesis, mechanism, and function. Cell 116, 281–297. 10.1016/s0092-8674(04)00045-5.

17. Pandit, K.V., Corcoran, D., Yousef, H., Yarlagadda, M., Tzouvelekis, A., Gibson, K.F., Konishi, K., Yousem, S.A., Singh, M., Handley, D., et al. (2010). Inhibition and role of let-7d in idiopathic pulmonary fibrosis. Am J Respir Crit Care Med 182, 220–229. 10.1164/rccm.200911-1698OC.

18. Cushing, L., Kuang, P.P., Qian, J., Shao, F., Wu, J., Little, F., Thannickal, V.J., Cardoso, W.V., and Lu, J. (2011). miR-29 is a major regulator of genes associated with pulmonary fibrosis. Am J Respir Cell Mol Biol 45, 287–294. 10.1165/rcmb.2010-0323OC.

19. Xiao, J., Meng, X.M., Huang, X.R., Chung, A.C., Feng, Y.L., Hui, D.S., Yu, C.M., Sung, J.J., and Lan, H.Y. (2012). miR-29 inhibits bleomycin-induced pulmonary fibrosis in mice. Mol Ther 20, 1251–1260. 10.1038/mt.2012.36.

20. Yan, L., Su, Y., Hsia, I., Xu, Y., Vincent-Chong, V.K., Mojica, W., Seshadri, M., Zhao, R., and Wu, Y. (2023). Delivery of anti-microRNA-21 by lung-targeted liposomes for pulmonary fibrosis treatment. Mol Ther Nucleic Acids 32, 36–47. 10.1016/j.omtn.2023.02.031.

21. Yamada, M., Kubo, H., Ota, C., Takahashi, T., Tando, Y., Suzuki, T., Fujino, N., Makiguchi, T., Takagi, K., Suzuki, T., and Ichinose, M. (2013). The increase of microRNA-21 during lung fibrosis and its contribution to epithelial-mesenchymal transition in pulmonary epithelial cells. Respir Res 14, 95. 10.1186/1465-9921-14-95.

22. Yang, S., Banerjee, S., de Freitas, A., Sanders, Y.Y., Ding, Q., Matalon, S., Thannickal, V.J., Abraham, E., and Liu, G. (2012). Participation of miR-200 in pulmonary fibrosis. Am J Pathol 180, 484–493. 10.1016/j.ajpath.2011.10.005.

23. Liang, H., Gu, Y., Li, T., Zhang, Y., Huangfu, L., Hu, M., Zhao, D., Chen, Y., Liu, S., Dong, Y., et al. (2014). Integrated analyses identify the involvement of microRNA-26a in epithelial-mesenchymal transition during idiopathic pulmonary fibrosis. Cell Death Dis 5, e1238. 10.1038/cddis.2014.207.

24. Yan, W., Wu, Q., Yao, W., Li, Y., Liu, Y., Yuan, J., Han, R., Yang, J., Ji, X., and Ni, C. (2017). MiR-503 modulates epithelial-mesenchymal transition in silica-induced pulmonary fibrosis by targeting PI3K p85 and is sponged by lncRNA MALAT1. Sci Rep 7, 11313. 10.1038/s41598-017-11904-8.

25. Kurowska-Stolarska, M., Hasoo, M.K., Welsh, D.J., Stewart, L., McIntyre, D., Morton, B.E., Johnstone, S., Miller, A.M., Asquith, D.L., Millar, N.L., et al. (2017). The role of microRNA-155/liver X receptor pathway in experimental and idiopathic pulmonary fibrosis. J Allergy Clin Immunol 139, 1946–1956. 10.1016/j.jaci.2016.09.021.

26. Wang, Y., Huang, C., Reddy Chintagari, N., Bhaskaran, M., Weng, T., Guo, Y., Xiao, X., and Liu, L. (2013). miR-375 regulates rat alveolar epithelial cell trans-differentiation by inhibiting Wnt/beta-catenin pathway. Nucleic Acids Res 41, 3833–3844. 10.1093/nar/gks1460.

27. Moimas, S., Salton, F., Kosmider, B., Ring, N., Volpe, M.C., Bahmed, K., Braga, L., Rehman, M., Vodret, S., Graziani, M.L., et al. (2019). miR-200 family members reduce senescence and restore idiopathic pulmonary fibrosis type II alveolar epithelial cell transdifferentiation. ERJ Open Res 5. 10.1183/23120541.00138-2019.

28. Volpe, M.C., Ciucci, G., Zandomenego, G., Vuerich, R., Ring, N.A.R., Vodret, S., Salton, F., Marchesan, P., Braga, L., Marcuzzo, T., et al. (2023). Flt1 produced by lung endothelial cells impairs ATII cell transdifferentiation and repair in pulmonary fibrosis. Cell Death Dis 14, 437. 10.1038/s41419-023-05962-2.

29. Chioccioli, M., Roy, S., Newell, R., Pestano, L., Dickinson, B., Rigby, K., Herazo-Maya, J., Jenkins, G., Ian, S., Saini, G., et al. (2022). A lung targeted miR-29 mimic as a therapy for pulmonary fibrosis. EBioMedicine 85, 104304. 10.1016/j.ebiom.2022.104304.

30. Wang, J.H., Gessler, D.J., Zhan, W., Gallagher, T.L., and Gao, G. (2024). Adeno-associated virus as a delivery vector for gene therapy of human diseases. Signal Transduct Target Ther 9, 78. 10.1038/s41392-024-01780-w.

31. Thomas, S.P., Domm, J.M., van Vloten, J.P., Xu, L., Vadivel, A., Yates, J.G.E., Pei, Y., Ingrao, J., van Lieshout, L.P., Jackson, S.R., et al. (2023). A promoterless AAV6.2FF-based lung gene editing platform for the correction of surfactant protein B deficiency. Mol Ther 31, 3457–3477. 10.1016/j.ymthe.2023.10.002.

32. van Lieshout, L.P., Domm, J.M., Rindler, T.N., Frost, K.L., Sorensen, D.L., Medina, S.J., Booth, S.A., Bridges, J.P., and Wootton, S.K. (2018). A Novel Triple-Mutant AAV6 Capsid Induces Rapid and Potent Transgene Expression in the Muscle and Respiratory Tract of Mice. Mol Ther Methods Clin Dev 9, 323–329. 10.1016/j.omtm.2018.04.005.

33. Rindler, T.N., Brown, K.M., Stockman, C.A., van Lieshout, L.P., Martin, E.P., Weaver, T.E., Zacharias, W.J., Wootton, S.K., Whitsett, J.A., and Bridges, J.P. (2021). Efficient Transduction of Alveolar Type 2 Cells with Adeno-associated Virus for the Study of Lung Regeneration. Am J Respir Cell Mol Biol 65, 118–121. 10.1165/rcmb.2021-0049LE.

34. Kang, M.H., van Lieshout, L.P., Xu, L., Domm, J.M., Vadivel, A., Renesme, L., Muhlfeld, C., Hurskainen, M., Mizikova, I., Pei, Y., et al. (2020). A lung tropic AAV vector improves survival in a mouse model of surfactant B deficiency. Nat Commun 11, 3929. 10.1038/s41467-020-17577-8.

35. Jenkins, R.G., Moore, B.B., Chambers, R.C., Eickelberg, O., Konigshoff, M., Kolb, M., Laurent, G.J., Nanthakumar, C.B., Olman, M.A., Pardo, A., et al. (2017). An Official American Thoracic Society Workshop Report: Use of Animal Models for the Preclinical Assessment of Potential Therapies for Pulmonary Fibrosis. Am J Respir Cell Mol Biol 56, 667–679. 10.1165/rcmb.2017-0096ST.

36. Liang, J., and Ligresti, G. (2023). Aging Delays Lung Repair: Insights from Omics Analysis in Mice with Pulmonary Fibrosis. Am J Respir Cell Mol Biol 69, 376–377. 10.1165/rcmb.2023-0171ED.

37. Wang, Y., Huang, C., Chintagari, N.R., Xi, D., Weng, T., and Liu, L. (2015). miR-124 regulates fetal pulmonary epithelial cell maturation. Am J Physiol Lung Cell Mol Physiol 309, L400–413. 10.1152/ajplung.00356.2014.

38. Zepp, J.A., and Morrisey, E.E. (2019). Cellular crosstalk in the development and regeneration of the respiratory system. Nat Rev Mol Cell Biol 20, 551–566. 10.1038/s41580-019-0141-3.

39. Sawhney, A.S., Deskin, B.J., Cai, J., Gibbard, D., Ali, G., Utoft, A., Qi, X., Olson, A., Hausman, H., Sabol, L., et al. (2025). A molecular circuit regulates fate plasticity in emerging and adult AT2 cells. Nat Commun 16, 8924. 10.1038/s41467-025-64224-1.

40. Le, H.Q., Hill, M.A., Kollak, I., Keck, M., Schroeder, V., Wirth, J., Skronska-Wasek, W., Schruf, E., Strobel, B., Stahl, H., et al. (2021). An EZH2-dependent transcriptional complex promotes aberrant epithelial remodelling after injury. EMBO Rep 22, e52785. 10.15252/embr.202152785.

41. Han, G., Sinjab, A., Rahal, Z., Lynch, A.M., Treekitkarnmongkol, W., Liu, Y., Serrano, A.G., Feng, J., Liang, K., Khan, K., et al. (2024). An atlas of epithelial cell states and plasticity in lung adenocarcinoma. Nature 627, 656–663. 10.1038/s41586-024-07113-9.

42. Li, J., and Liu, L. (2025). miR-124-3p inhibits CRC proliferation, migration, and invasion by targeting ITGB1. Discov Oncol 16, 158. 10.1007/s12672-025-01936-2.

43. Dong, Z.B., Wu, H.M., He, Y.C., Huang, Z.T., Weng, Y.H., Li, H., Liang, C., Yu, W.M., and Chen, W. (2022). MiRNA-124-3p.1 sensitizes hepatocellular carcinoma cells to sorafenib by regulating FOXO3a by targeting AKT2 and SIRT1. Cell Death Dis 13, 35. 10.1038/s41419-021-04491-0.

44. Ma, T., Zhao, Y., Wei, K., Yao, G., Pan, C., Liu, B., Xia, Y., He, Z., Qi, X., Li, Z., et al. (2016). MicroRNA-124 Functions as a Tumor Suppressor by Regulating CDH2 and Epithelial-Mesenchymal Transition in Non-Small Cell Lung Cancer. Cell Physiol Biochem 38, 1563–1574. 10.1159/000443097.

45. Ji, H., Sang, M., Liu, F., Ai, N., and Geng, C. (2019). miR-124 regulates EMT based on ZEB2 target to inhibit invasion and metastasis in triple-negative breast cancer. Pathol Res Pract 215, 697–704. 10.1016/j.prp.2018.12.039.

46. Qian, W., Cai, X., Qian, Q., Peng, W., Yu, J., Zhang, X., Tian, L., and Wang, C. (2019). lncRNA ZEB1-AS1 promotes pulmonary fibrosis through ZEB1-mediated epithelial-mesenchymal transition by competitively binding miR-141-3p. Cell Death Dis 10, 129. 10.1038/s41419-019-1339-1.

47. Chilosi, M., Calio, A., Rossi, A., Gilioli, E., Pedica, F., Montagna, L., Pedron, S., Confalonieri, M., Doglioni, C., Ziesche, R., et al. (2017). Epithelial to mesenchymal transition-related proteins ZEB1, beta-catenin, and beta-tubulin-III in idiopathic pulmonary fibrosis. Mod Pathol 30, 26–38. 10.1038/modpathol.2016.147.

48. Yao, L., Conforti, F., Hill, C., Bell, J., Drawater, L., Li, J., Liu, D., Xiong, H., Alzetani, A., Chee, S.J., et al. (2019). Paracrine signalling during ZEB1-mediated epithelial-mesenchymal transition augments local myofibroblast differentiation in lung fibrosis. Cell Death Differ 26, 943–957. 10.1038/s41418-018-0175-7.

49. Kobayashi, H., Tachi, A., and Hagita, S. (2024). Time course of histopathology of bleomycin-induced pulmonary fibrosis using an intratracheal sprayer in mice. Exp Anim 73, 41–49. 10.1538/expanim.23-0048.

50. Brazee, P., Allen, N., Knipe, R., Redente, E.F., and Le Saux, C.J. (2025). Peeling Back the Layers of the Bleomycin Model of Lung Fibrosis: Lessons Learned, Factors to Consider, and Future Directions. Semin Respir Crit Care Med 46, 330–346. 10.1055/a-2649-9402.

51. Desai, T.J., Brownfield, D.G., and Krasnow, M.A. (2014). Alveolar progenitor and stem cells in lung development, renewal and cancer. Nature 507, 190–194. 10.1038/nature12930.

52. Neo, W.H., Yap, K., Lee, S.H., Looi, L.S., Khandelia, P., Neo, S.X., Makeyev, E.V., and Su, I.H. (2014). MicroRNA miR-124 controls the choice between neuronal and astrocyte differentiation by fine-tuning Ezh2 expression. J Biol Chem 289, 20788–20801. 10.1074/jbc.M113.525493.

53. Ponomarev, E.D., Veremeyko, T., Barteneva, N., Krichevsky, A.M., and Weiner, H.L. (2011). MicroRNA-124 promotes microglia quiescence and suppresses EAE by deactivating macrophages via the C/EBP-alpha-PU.1 pathway. Nat Med 17, 64–70. 10.1038/nm.2266.

54. Reyfman, P.A., Walter, J.M., Joshi, N., Anekalla, K.R., McQuattie-Pimentel, A.C., Chiu, S., Fernandez, R., Akbarpour, M., Chen, C.I., Ren, Z., et al. (2019). Single-Cell Transcriptomic Analysis of Human Lung Provides Insights into the Pathobiology of Pulmonary Fibrosis. Am J Respir Crit Care Med 199, 1517–1536. 10.1164/rccm.201712-2410OC.

55. Yao, C., Guan, X., Carraro, G., Parimon, T., Liu, X., Huang, G., Mulay, A., Soukiasian, H.J., David, G., Weigt, S.S., et al. (2021). Senescence of Alveolar Type 2 Cells Drives Progressive Pulmonary Fibrosis. Am J Respir Crit Care Med 203, 707–717. 10.1164/rccm.202004-1274OC.

56. Morse, C., Tabib, T., Sembrat, J., Buschur, K.L., Bittar, H.T., Valenzi, E., Jiang, Y., Kass, D.J., Gibson, K., Chen, W., et al. (2019). Proliferating SPP1/MERTK-expressing macrophages in idiopathic pulmonary fibrosis. Eur Respir J 54. 10.1183/13993003.02441-2018.

57. Wang, J., Qing, B., Gu, L., Chen, H., Chen, Y., Tang, Y., Ge, Z., Hu, R., Yuan, Y., and Xia, Z. (2025). Caspase-9 activates beta-catenin signaling to promote pulmonary fibrosis. J Transl Med 23, 986. 10.1186/s12967-025-07020-1.

58. Haak, A.J., Ducharme, M.T., Diaz Espinosa, A.M., and Tschumperlin, D.J. (2020). Targeting GPCR Signaling for Idiopathic Pulmonary Fibrosis Therapies. Trends Pharmacol Sci 41, 172–182. 10.1016/j.tips.2019.12.008.

59. Fortier, S.M., Redente, E.F., and Peters-Golden, M. (2025). Reimagining Fibrosis Research, Outcomes, and Therapeutics Through the Lens of Resolution. Semin Respir Crit Care Med 46, 298–310. 10.1055/a-2666-7479.

60. Maher, T.M., Assassi, S., Azuma, A., Cottin, V., Hoffmann-Vold, A.M., Kreuter, M., Oldham, J.M., Richeldi, L., Valenzuela, C., Wijsenbeek, M.S., et al. (2025). Nerandomilast in Patients with Progressive Pulmonary Fibrosis. N Engl J Med 392, 2203–2214. 10.1056/NEJMoa2503643.

61. Ibrahim, M., Piazza, G.A., and Ahsan, F. (2026). Nerandomilast as the first PDE4B-selective therapy in idiopathic pulmonary fibrosis. Trends Pharmacol Sci 47, 120–121. 10.1016/j.tips.2025.11.006.

62. Ciucci, G., Braga, L., and Zacchigna, S. (2025). Discovery platforms for RNA therapeutics. Br J Pharmacol 182, 281–295. 10.1111/bph.16424.

